# Molecular architecture of the mouse nervous system

**DOI:** 10.1101/294918

**Authors:** Amit Zeisel, Hannah Hochgerner, Peter Lönnerberg, Anna Johnsson, Fatima Memic, Job van der Zwan, Martin Häring, Emelie Braun, Lars Borm, Gioele La Manno, Simone Codeluppi, Alessandro Furlan, Nathan Skene, Kenneth D. Harris, Jens Hjerling Leffler, Ernest Arenas, Patrik Ernfors, Ulrika Marklund, Sten Linnarsson

## Abstract

The mammalian nervous system executes complex behaviors controlled by specialised, precisely positioned and interacting cell types. Here, we used RNA sequencing of half a million single cells to create a detailed census of cell types in the mouse nervous system. We mapped cell types spatially and derived a hierarchical, data-driven taxonomy. Neurons were the most diverse, and were grouped by developmental anatomical units, and by the expression of neurotransmitters and neuropeptides. Neuronal diversity was driven by genes encoding cell identity, synaptic connectivity, neurotransmission and membrane conductance. We discovered several distinct, regionally restricted, astrocytes types, which obeyed developmental boundaries and correlated with the spatial distribution of key glutamate and glycine neurotransmitters. In contrast, oligodendrocytes showed a loss of regional identity, followed by a secondary diversification. The resource presented here lays a solid foundation for understanding the molecular architecture of the mammalian nervous system, and enables genetic manipulation of specific cell types.

---

DESPITE A CENTURY OF EFFORTS using morphological, electrophysiological, histochemical and molecular approaches, we still lack systematic, comprehensive and detailed information about cell types in the nervous system. We do not know the number and variety of neurons, glia or vascular cells; we do not know their molecular relationships or local heterogeneity, and we have little systematic knowledge of the overall organization of the nervous system in molecular terms. Neurons are clearly highly diverse, but how much diversity is there among astrocytes, oligodendrocytes or vascular cells? Does the glial diversity reflect local environment (e.g. local neuronal types), developmental compartment, or some aspect of glia function?

The organization of the adult mammalian nervous system is the result of developmental, functional, evolutionary and biomechanial constraints. Our current understanding of its architecture originated with the pioneering studies of Santiago Ramón y Cajal, who mapped microscopic neuroanatomy in exquisite detail. The adult brain is organized into dorsoventral and rostrocaudal compartments, which result from patterning of the early neural tube (Rubenstein and Rakic, 2013). However, many neurons (e.g. telencephalic interneurons) and glia (e.g. oligodendrocyte precursor cells), vascular and immune cells migrate long distances during embryogenesis and thus end up in a location different from their place of birth. Furthermore, convergent functional specialization occurs in many parts of the nervous system: for example, dopaminergic neurons are found both in the midbrain and in the olfactory bulb, and noradrenergic neurons in the sympathetic ganglia as well as the hindbrain.

The question therefore arises if the molecular identity of a cell is determined mainly by its developmental ancestry, by its local environment, or by its function. All three possibilities are plausible a priori: neurons with shared function (for example, long-range projecting neurons, or neurons using a common principal neurotransmitter) might be expected show common gene expression states across brain regions. Alternatively, chemical cues arising from a local environment might impose constraints forcing neighboring cells of different functions to become molecularly similar. Finally, developmental origin, through shared gene regulatory circuits, might retain an imprint on cell types in the adult, so that gene expression patterns would reflect developmental domains and borders.

Recently, single-cell RNA sequencing (scRNAseq) has emerged as a powerful method for unbiased discovery of cell types and states (Islam et al., 2011, 2013; Jaitin et al., 2014; Macosko et al., 2015; Shekhar et al., 2016; Tang et al., 2009; Tasic et al., 2016; Usoskin et al., 2014; Zeisel et al., 2015), and initiatives are underway to create atlases of both human and model organisms (Han et al., 2018; Regev et al., 2017). Here, we used systematic scRNA-seq to survey cells across the central and peripheral nervous system. We use the inferred molecular relationships between all cell types to propose a data-driven taxonomy of cell types, and we discuss the overall architecture of the mammalian nervous system in light of this taxonomy.

## A molecular survey of the mouse nervous system

We performed a comprehensive survey of the adolescent mouse nervous system by single-cell RNA sequencing. We dissected the brain and spinal cord into contiguous anatomical regions, and further included the peripheral sensory, enteric and sympathetic nervous system. In total, we analyzed 19 regions (Figure 1A), but omitting at least the retina, the olfactory epithelium, the vomeronasal organ, the inner ear, and the parasympathetic ganglia.

**Figure 1.**
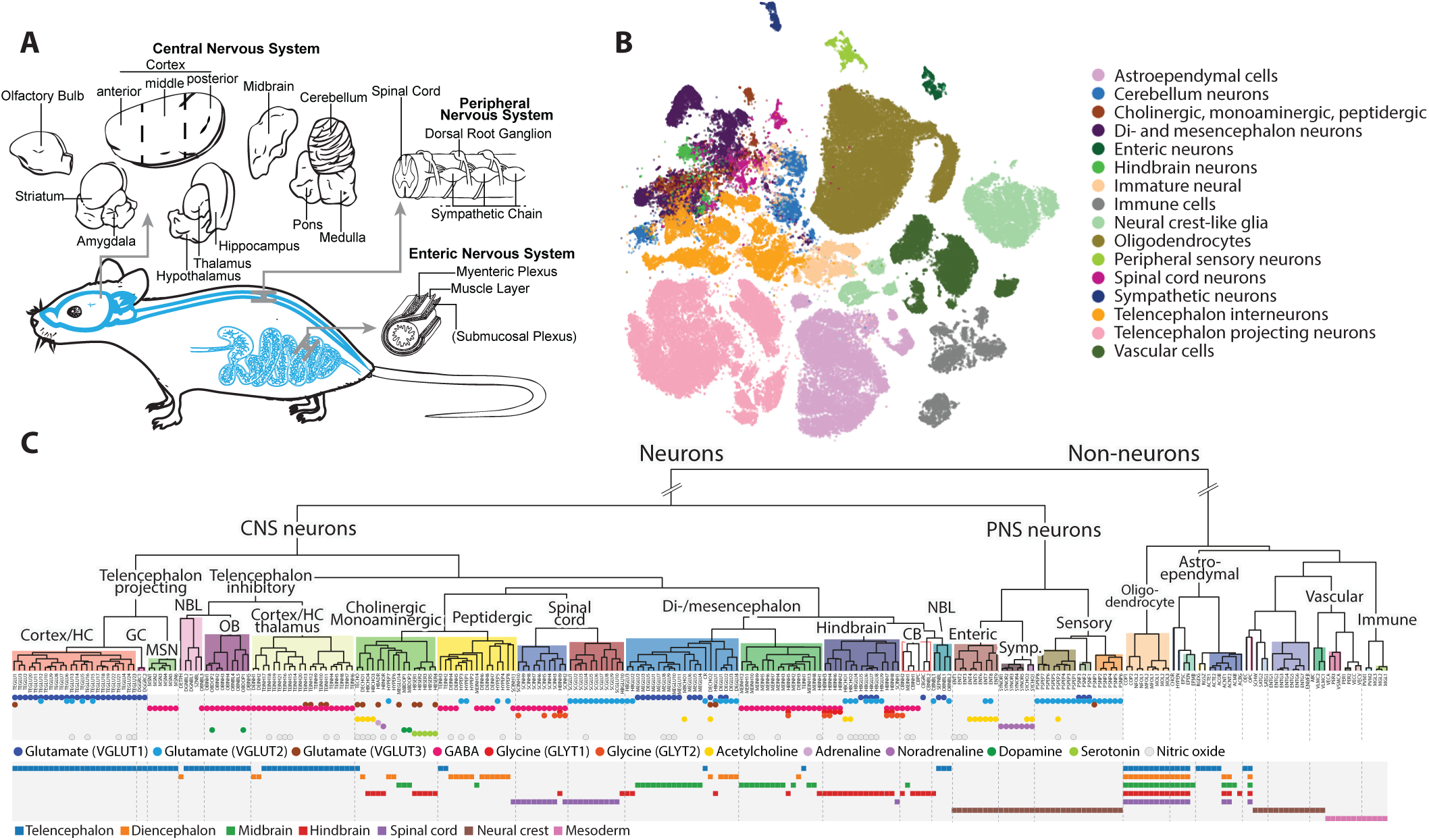
Molecular survey of the mouse nervous system using single-cell RNA sequencing. (A) Schematic illustation of the sampling strategy. The brain was divided into coarse anatomical units and in addition we sampled from the spinal cord, dorsal root ganglia, sympathetic ganglion and the enteric nervous system. (B) Visualization of the single cell data using gt-SNE embedding (see Methods). Cells are colored by rank 3 taxonomy units, indicated in the legend. (C) Dendrogram describing the taxonomy of all identified cell types. Main branches, corresponding to the taxonomy, are annotated with labels and colored background. The neurotransmitter used by each cell types is indicated below the leafs as colored circles. Lower panel indicate the developmental compartment of origin for each cell types.

For each region, we typically sampled from all cells without selection. However, in the enteric nervous system, we used *Wnt1-Cre;R26Tomato* transgenic mice to isolate myenteric plexus cells of the small intestine by FACS (we did not include the submucosal plexus or any other regions of the gastrointestinal tract). In the sympathetic nervous system, we used the same mice to guide dissection of the superior cervical and stellate ganglia, as well as thoracic ganglia 1-13 (we did not include lumbar ganglia). In the hippocampus and cortex we similarly isolated inhibitory neurons from the *vGat-Cre;TdTomato* strain by FACS. We used at least two mice for each tissue, typically one male and one female, and analyzed a total of 133 samples (Table S1) by droplet microfluidics (10X Genomics Chromium) to reveal the transcriptomes of 509,876 cells.

Preliminary analyses showed that the dataset contained hundreds of distinct cell types, and that the dynamic range of cell type abundances spanned four orders of magnitude. In addition, the dataset was affected by a number of technical artefacts, including low-quality cells, batch effects, sex-specific gene expression, neuronal activity-dependent gene expression, and more. To overcome these challenges, we developed a multistage analysis pipeline called cytograph, which progressively discovers cell types or states, while mitigating the impact of technical artefacts. For scalability, cytograph uses algorithms that scale approximately linearly with the number of cells, and automatically parallelizes tasks when possible (Methods).

After an initial quality assessment of samples and cells, we retained 492,949 cells as inputs to the computational analysis. During three stages of manifold learning and clustering we removed additional doublets, outliers and low-quality cells (Fig. S1A). As oligodendrocytes are extremelyabundant in the hindbrain and spinal cord, we removed more than 200,000 oligodendrocytes from these regions, in order to better balance the number of oligodendrocytes between tissues (but analyzing the full set of cells did not reveal any additional structure in the oligodendrocyte lineage). We then manually curated the automatically generated clusters, removing additional low-quality clusters and merging highly similar clusters. The final, high-quality curated compendium comprised 265 clusters represented by 160,796 high-quality single-cell transcriptomes (Fig. 1B-C). This represents a highly conservative clustering, and significant heterogeneity likely remains within many of the reported clusters.

To assess the robustness of the clusters, we trained a random forest classifier to recognize cluster labels and then assessed its performance on held-out data (80% training set, 20% test set). The average precision and recall were both 82%, indicating a high level of robustness, particularly considering the large total number of clusters. Most classification errors occurred between closely related cell types. To reveal this type of relatedness of clusters, we computed the probability, for each cluster, that its cells would be classified as any other type (Fig. S1D). Mostly, only the correct cluster showed high probability, but in other cases the probability distribution revealed relatedness between sets of biologically related clusters (e.g. hindbrain serotonergic neurons).

In order to validate our ability to recover known cell types, we next assessed the concordance with six different previously published and experimentally validated scRNA-seq datasets, comprising two different technologies (Fludigm C1 and 10X Genomics Chromium) and five tissues: cortex (Zeisel et al., 2015), striatum (Muñoz-Manchado, in preparation), dentate gyrus (Hochgerner et al., 2018), spinal cord (Häring et al. 2018, in press) and sympathetic nervous system (Furlan et al., 2016). Of the 139 previously published clusters, 98% were perfect or near-perfect matches to corresponding clusters in the new compendium (84% perfect, 14% near-perfect, 2% mismatches; see Table S2).

We performed a comprehensive annotation of the clusters using a variety of automated and manual methods. We assigned each cluster a unique mnemonic identifier (e.g. MBDOP1), a descriptive name (“Midbrain dopaminergic neuron”), a major class (e.g. “neuron”), neurotransmitter identity, putative developmental origin, anatomical location and region (Table S3).

To characterize gene expression across clusters, we computed enriched genes for each cluster, indicating increased but not unique expression. We also computed a probabilistic “trinarization” score, which can be used to determine if a gene is expressed, not expressed, or ambiguous, in each cluster (Methods). We combined enrichment and trinarization scores to discover marker gene sets sufficient to uniquely identify each cluster, with high probability. Remarkably, we found that 248 (93%) of all clusters were uniquely identifiable with just two genes, while 17 required three genes, and none required more than three (although, adding more genes could increase the robustness of identification). This finding attests to the precise molecular organization of the mammalian nervous system, and shows that nearly all cell types can be genetically accessed and manipulated using readily available intersectional gene targeting approaches (Allen and Luo, 2015).

### Box 1 Resources

The raw sequence data is deposited in the sequence read archive under accession SRP135960.

The companion wiki at http://mousebrain.org, provides a report card for each cell type. The wiki can be browsed by taxon, cell type, tissue, and gene, with information on enriched genes, specific markers, anatomical location and more. The download section of the wiki makes available the following resources:

- Aligned reads in the form of BAM files.
- Quality-control results of each sample (10X Genomics *cellranger* QC output).
- Expression data organized by individual Chromium sample, region, taxonomic group, and the entire final curated dataset. These files contain full metadata, graph layout, cluster assignments and cell type/state annotations, where appropriate.

Expression data is provided in Loom format (see http://loompy.org) and comes with an interactive, web-based viewer for explorative analysis. The wiki provides links to relevant Loom files, preloaded in the Loom viewer.

The analysis software developed for this paper is available at https://github.com/linnarsson-lab, in repositories named *cytograph* and *adolescent-mouse*.

We trained a support-vector machine classifier to automatically assign each cell to one of seven major classes: neurons, oligodendrocytes (all ~236,000), astrocytes, ependymal cells, peripheral glia (e.g. Schwann cells, satellite and enteric glia), immune cells and vascular cells (Fig. S1B). Neurons were most prevalent in rostral regions of the central nervous system (CNS), as well as in the cerebellum. In telencephalon, 61% of cells were neurons, compared with 43% in diencephalon, 19% in midbrain and 6% in hindbrain and spinal cord (not including cerebellum) and 78% in the cerebellum. In caudal regions, oligodendrocytes—needed to support long-range neurotransmission—dominated greatly, comprising 84% of cells in the hindbrain (excepting cerebellum) and 71% in the spinal cord. Astrocytes ranged from 13% of cells in the telencephalon to 6% in the hindbrain. In the peripheral nervous system (PNS), neurons were again the most common class in sensory and sympathetic ganglia (62% and 87%, respectively), whereas the enteric nervous system comprised 91% enteric glia and only 7% neurons. Due to sources of bias such as differential survival or cell capture, these estimates can be only approximate, but they are in good agreement with previous findings using scRNA-seq (Zeisel et al., 2015).

For a detailed description of all experimental and computational procedures, see Methods. An interactive browser, and links to all the data, code and annotations, is provided as a companion web site (see Box 1).

TO BEGIN TO UNDERSTAND the molecular organization of the mammalian nervous system, we calculated a robust dendrogram of cell types (Fig. 1C, Methods), showing relationships between cell types based on gene expression distance. The resulting arrangement of cell types revealed a surprisingly simple organization of the mammalian nervous system, which was largely organized according to three overlapping and interacting principles: major class (e.g neurons, astrocytes), developmental origin (e.g. telencephalon, diencephalon, midbrain, hindbrain) and neurotransmitter type (e.g. GABA, glutamate).

At the top level, neurons were separated from non-neuronal types regardless of tissue, reflecting a split between major classes of cells that express thousands of genes differentially. Notably, this first split does not correspond to any shared developmental or anatomical origins, as it groups neurons from both the central and peripheral nervous systems on one side, and the corresponding central (e.g. astrocytes) and peripheral (e.g. Schwann cells) glia on the other, along with developmentally unrelated vascular and immune cells.

The second level divided neurons according to their major region of origin. PNS neurons segregated from the CNS, reflecting the developmental split between neural crest-derived (PNS) and neural tube-derived (CNS) neurons. The peripheral neurons then split into sensory, sympathetic and enteric subdivisions, corresponding to both functional, anatomical and developmental differences between the three major divisions of the peripheral nervous system.

CNS neurons generally split first by anteroposterior domain (olfactory, telencephalon, diencephalon, midbrain, hindbrain, spinal cord), and then by excitatory versus inhibitory neurotransmitter. However, there were interesting exceptions, which will be explored further below.

Based on these and similar observations, we propose a data-driven molecular taxonomy arranged in a hierarchy of more than 70 named taxa (Table 1 and Figs. S2 and S3), respecting the dendrogram of Figure 1. The taxonomy provides an objective structuring principle for exploring the global architecture of the mammalian nervous system. Note that, where the taxonomy conflicted with properties of a cell type, we nevertheless assigned it to a taxon according to the dendrogram. For example, a spinal cord cell type that ended up in the hindbrain part of the dendrogram was assigned to the *Hindbrain* taxon (but was still individually named and annotated according to its true location, e.g. SCINH1, spinal cord inhibitory neurons).

## Postnatal neurogenesis in the CNS

Although most neuronal types were already mature at the age investigated (postnatal day 20-30), we observed signs of ongoing neurogenesis in several regions (Fig. 2). As expected, we detected the two regions that maintain adult neurogenesis in the mouse: the subventricular zone along the striatum and the dentate gyrus subgranular zone. In the subventricular zone, radial glia-like cells (RGSZ) and cycling neuronal intermediate progenitor cells (SZNBL) were linked to more mature and presumably migrating neuroblasts along the rostral migratory stream and in the olfactory bulb (OBNBL3). In the subgranular zone of the dentate gyrus, radial glia-like cells (RGDG), neuroblasts (NBDG) and immature granule cells (DGNBL1 and DGNBL2) would give rise to mature granule cells (DGGC), as recently described in detail (Hochgerner et al., 2018). The radial glia-like cells (RGSZ and RGDG), which are the stem cells of both lineages, were closely related and were more similar to astrocytes than to any neuroblast. They expressed *Riiad1* (shared with ependymal cells; Fig 2D), a gene of uncertain function otherwise found in lung, testis and the adrenal gland. We speculate that this gene is functionally required by ciliated cells such as radial glia, ependymal cells, sperm, lung ciliated epithelium and adrenal chromaffin cells which are common to these tissues. However, within the radial glia we found that RGDG and RGSZ further presented specific markers representing the local neurogenic niche (Fig. S4; for example the transcription factor *Tfap2c* in RGDG and *Urah* in RGSZ).

**Figure 2.**
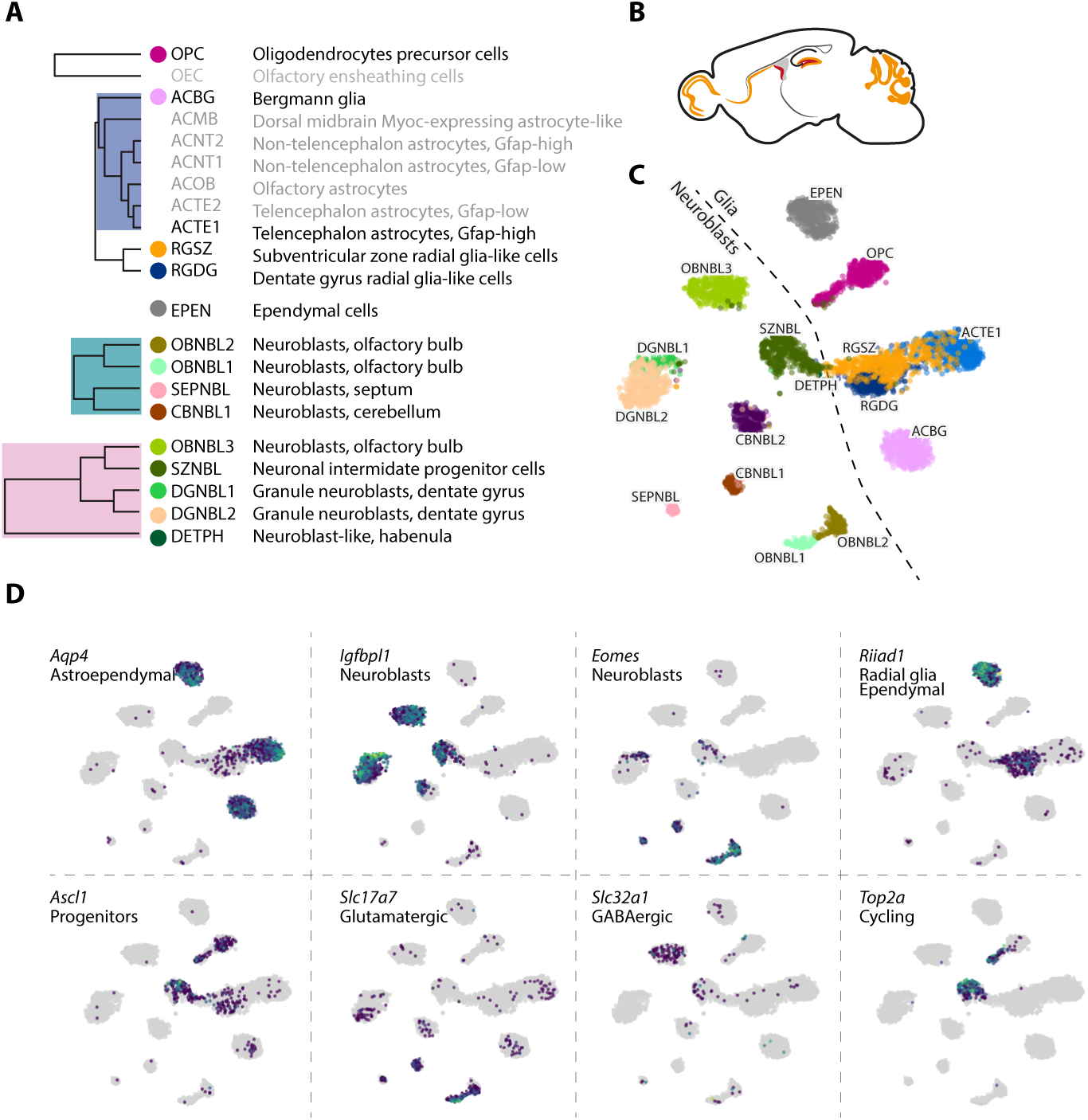
A map of neurogenesis in the juvenile mouse brain. (A) A cut-out from dendrogram of relevant cell types including neuroblast, radial glia-like, astrocytes, OPC and ependymal cells. (B) Sketch illustrating the locations where we found neurogenic activity. (C) gt-SNE embedding of all cells from the relevant cell types shown in A. Dashed line suggests the border between glia-like cells and neuroblasts. (D) Expression distribution of individual key genes projected onto the gt-SNE embedding.

Neuroblasts across the brain fell into two general categories, represented by two subtrees in the dendrogram (labelled NBL in Fig1D). The first category expressed *Igfbpl1* and was either GABAergic (OBNBL3) or did not express any clear neurotransmitter phenotype. These neuroblasts were found in the rostral migratory stream (SZNBL and OBNBL3), dentate gyrus (DGNBL2) and in the habenula (DETPH). The second category expressed the T-box transcription factor *Eomes* (also known as *Tbr2*) and the vesicular glutamate transporter (*Slc17a7*, also known as VGLUT1), hence was glutamatergic. These neuroblasts were found in the olfactory bulb (OBNBL1, OBNBL2), cerebellum (CBNBL) and septum (SEPNBL). However, *Eomes* and *Igfbpl1* overlapped in some populations (DGNBL1 and to some extent SZNBL), indicating that these categories of neuroblasts may represent sequential stages of neuronal maturation, rather than divergent cell types. *Eomes*-expressing neuroblasts, with a generally less mature neurotransmitter phenotype, may then represent early stages of neuronal differentiation, whereas *Igfbpl1* spans both early and later stages, as was already shown in the dentate gyrus (Hochgerner et al., 2018). Our data suggest that the presence of neurogenesis in the juvenile brain is more widespread than previously appreciated. Although late neurogenesis in the cerebellum, olfactory bulb, the rostral migratory stream, the dentate gyrus and the hippocampus have been well studied, importantly we now provide distinct molecular markers of all these populations or stages from RGL to neuroblasts (Figure S4). Furthermore, we identify immature neuronal cell types in the septum and habenula, which have previously been only poorly described.

## Astroependymal cells are diverse and spatially patterned

Astrocytes, ependymal cells and radial glia are developmentally related cell types, and formed a subtree in the dendrogram (Fig. 3). This taxon included two specialized secretory cell types: the hypendymal cells, HYPEN, which are specialized ependymal-like cells of the subcommissural organ that secrete SCO-spondin (encoded by *Sspo*) into the cerebrospinal fluid to form Reissner’s fiber; and the choroid plexus epithelial cells, CHOR, which are an extension of the ependymal lining of the ventricular surfaces that envelop branching capillaries protruding into the ventricles, and secrete the extremely abundant thyroxine and retinol transport protein Transthyretin (encoded by *Ttr*).

**Figure 3.**
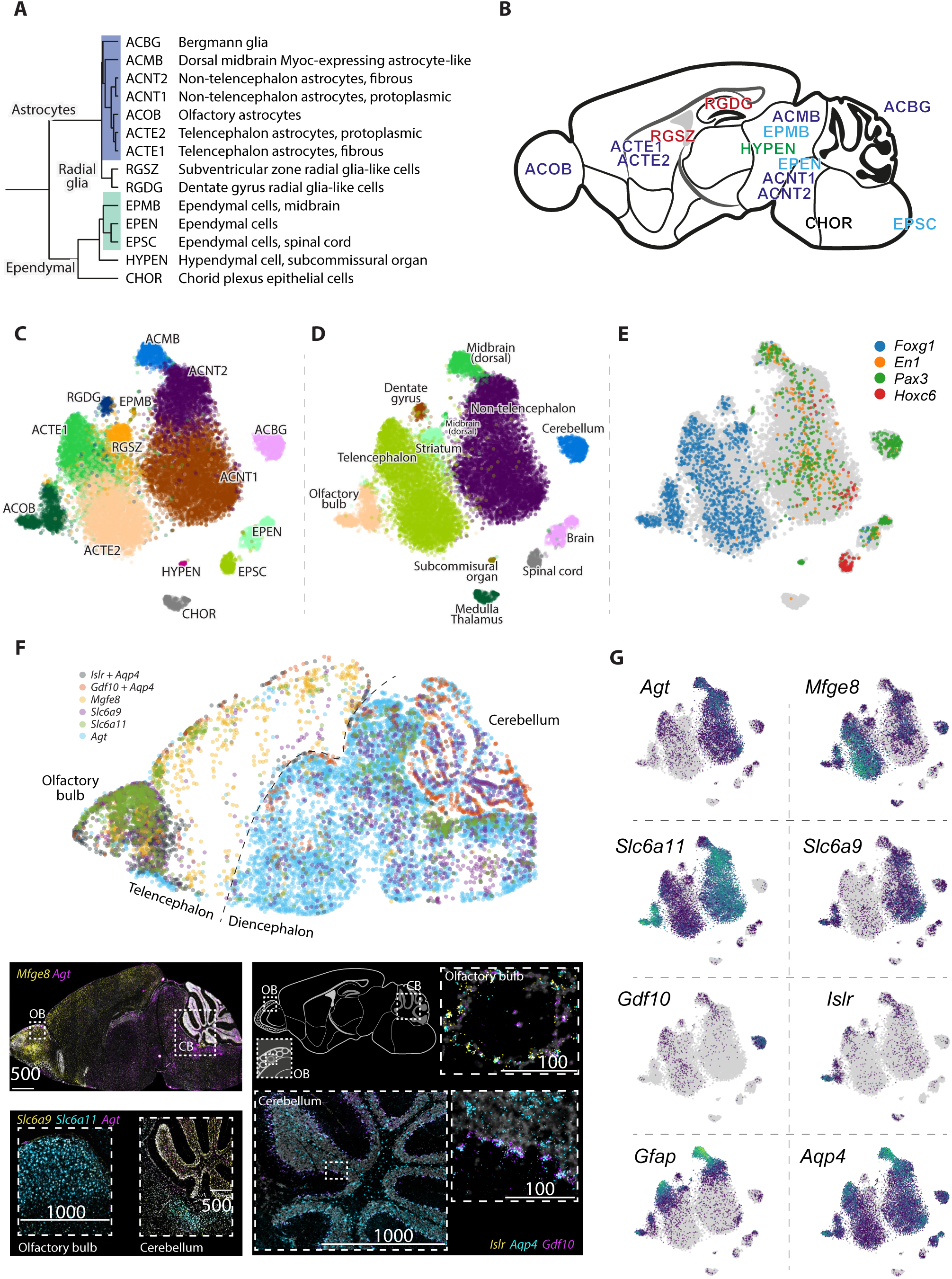
Molecular and spatial diversity of the astroependymal cells in the CNS. (A) Subtree describing the hierarchy of astroependymal cell types. (B) Schematic sagittal section showing the location of astroependymal cells. (C-E) gt-SNE embedding of all cells from the relevant clusters colored by cluster identity (C), tissue of origin (D) and patterning transcription factors (E). (F) Validation of spatial distribution of astrocytes cell types using multiplex in-situ hybridization (RNAscope). Images from three consecutive sections were aligned and overlaid (see Methods) to generate a composite with dots representing cells (upper panel). Below, high-magnification images showing details of spatial location. (G) Gene expression of selected markers shown on the gt-SNE layout.

Three types of ependymal all expressed *Foxj1*, the master regulator of motile cilia (Yu et al., 2008). The first, EPEN, was common along the rostrocaudal axis. The second, EPMB, was observed in the dorsal midbrain and—to a lesser extent—the hypothalamus. They expressed high levels of *Gfap* and the *Efnb3* gene encoding Ephrin B3, but only low levels of *Foxj1*. They also expressed many markers of tanycytes of the third ventricle, including *Nes*, *Vim*, *Rax* and *Gpr50* (Miranda-Angulo et al., 2014), but their location in the dorsal midbrain suggests that they instead represented a tanycyte of the circumventricular organs (Kettenmann and Ransom, 2013). The third, EPSC, was specific to the spinal cord and was distinguished by the expression of immediate-early genes such as *Fos*, *Junb* and *Egr1*.

ASTROCYTES WERE DESCRIBED in 1856 by Rudolf Virchow as nervenkitt—neuroglia—and in the second half of the 19th century a number of distinct astrocyte types were identified (Somjen, 1988). Specialized retinal and cerebellar radial glia were described by Heinrich Müller (1851) and Karl Bergmann (1857). Common stellate astrocytes were described by Otto Deiters in 1865, studied in detail by Camillo Golgi in the 1870s, and classified into protoplasmic (grey matter) and fibrous (white matter) by 1893. With the exception of the discovery of radial astrocytes in neurogenic regions of the brain (see above), reactive astrocytes in response to injury, and types defined solely by morphology such as velate astrocytes of the cerebellum and olfactory bulb, the modern understanding of astrocyte diversity essentially stands as it stood in 1900: the main types of mature astrocytes are believed to be the Müller glia, the Bergmann glia, and the protoplasmic and fibrous astrocytes (Ben Haim and Rowitch, 2016).

Here, we observed seven molecularly distinct types of astrocytes with clear regionally specialized distribution. All astrocytes expressed *Aqp4*, encoding aquaporin 4, the water channel located on astrocyte vascular end feet. In addition to Bergmann glia of the cerebellum (ACBG), we found olfactory-specific astrocytes (ACOB, unrelated to olfactory ensheathing cells; see below), two subtypes of telencephalon-specific astrocytes (ACTE1 and ACTE2), two subtypes of non-telencephalon astrocytes (ACNT1 and ACNT2), and a *Myoc*-expressing astrocyte of the dorsal midbrain, ACMB. Müller glia were not observed because we did not sample from the retina.

Olfactory astrocytes were located around the olfactory glomeruli, and could represent the previously described velate astrocytes, for which no molecular properties are known; they highly specifically expressed the *Islr* and *Islr2* genes encoding immunoglobulin-domain cell adhesion proteins, (Figure 3, Figure S5) among other genes.

Telencephalon astrocytes ACTE1 and ACTE2 were distinguished by the expression of several genes including *Mfge8*, *Lhx2* and were found in the olfactory bulb, cerebral cortex, striatum, amygdala and hippocampus, but absent from hypothalamus, thalamus, midbrain and hindbrain. Non-telencephalic astrocytes ACNT1 and ACNT2 showed the opposite distribution, marked by *Agt* (angiotensinogen) and found in all regions caudal to the telencephalon/diencephalon border (i.e. posterior to and including the hypothalamus and thalmus). The border between the two was sharp, as judged by in situ hybridization of the relevant genes (Fig. S5C), indicating that they do not intermingle across substantial distances.

We validated the identity and distribution of astrocyte cell types using RNA FISH (RNAscope) which was fully consistent with *in situ* hybridization (Figs. 3F and S5). Co-staining of *Mfge8* and *Agt* on a sagittal section revealed a clear border separating the telencephalon from the diencephalon. Olfactory bulb and cerebellum were enriched with their local astrocytes ACOB and ACBG marked by *Islr* and *Gdf10* respectively. Moreover, we validated the distribution of neurotransmitter transporters with *Slc6a11* (also known as GAT3, the GABA reuptake transporter) highly expressed in the olfactory and the non-telencephalon astrocytes (but not in cerebellum) and *Slc6a9* (glycine transporter GLYT1) with similar pattern but lower olfactory expression and a higher expression in the cerebellum.

Both telencephalon and non-telencephalon astrocytes were further split into subtypes expressing *Gfap* at high or low levels (Fig. 3G). This distinction likely corresponds to the fibrous astrocytes of the white matter and the glia limitans underneath the pia (*Gfap*-high) versus the protoplasmic astrocytes of the parenchyma (*Gfap*-low). The difference between subtypes in both cases involved a similar set of genes, suggesting that this represents an independent axis of variation that can be activated in both telencephalon and non-telencephalon astrocytes as a function of local environmental cues, particularly the distance from the pia and white matter.

Interestingly, like neurons, these diverse astrocyte and ependymal cell types occupied distinct domains of the brain with little apparent mixing. The sharpness of the border between *Mfge8* (telencephalon astrocytes) and *Agt* (non-telencephalon) expression, for example, and the fact that it coincided with a developmentally recognized boundary distinguishing the telencephalon from the rest of the brain, strongly implies that these astrocyte types are developmentally specified. In order to test this hypothesis, we examined the expression of region-specific neural tube patterning genes, the transcription factors *Foxg1* (telencephalon), *En1* and *Pax3* (midbrain) and *Hoxc6* (spinal cord). Each of these genes marked the expected subset of astrocyte and ependymal cell types (Fig. 3G). For example, *Foxg1* labelled ACOB, ACTE1 and ACTE2 as well as radial glia of the striatum and dentate gyrus (RGSZ and RGDG), but not the non-telencephalon astrocytes or any of the specialized ependymal and choroid cells (e.g. ACMB, EPMB, CHOR, HYPEN, and EPSC). Conversely, *Hoxc6* labelled a subset of the non-telencephalic astrocytes as well as spinal cord ependymal cells. The common, brain-wide ependymal cells (EPEN) were labelled by both *Foxg1*, *En1* and *Pax3*, but not by *Hoxc6*, in agreement with their brain-wide distribution. Thus we have uncovered a previously unrecognized diversity of astrocyte and ependymal cell types, showing the hallmarks of developmentally specified identities and regional specialization.

We can only speculate as to the functional distinction between telencephalic and non-telencephalic astrocytes. Given the important role of astrocytes in maintaining neurotransmission, it’s striking that the distinction between telencephalic and non-telencephalic astrocytes coincided with the prevalence of VGLUT1 in the telencephalon versus VGLUT2 in the di-/mesencephalon and hindbrain (Fig. S1C and S5F; however, the thalamus used both VGLUTs). This indicates a possible role in maintaining distinct modes of glutamatergic neurotransmission.

Furthermore, one of the genes most highly enriched in non-telencephalic astrocytes was Slc6a9 (Fig. 3G), encoding the glycine reuptake transporter GLYT1. Glycine is a widely used inhibitory neurotransmitter only in the caudal parts of the brain and in the spinal cord. This suggests a specific role for non-telencephalic astrocytes in clearing glycine from the synaptic cleft. We note that since GLYT1 is a reversible glycine transporter (Supplisson and Roux, 2002), if it were expressed in telencephalic astrocytes then those cells would potentially secrete glycine into the synaptic cleft instead of absorbing it. Glycine is not only an inhibitory neu-rotransmitter, but also an co-ligand for the NMDA glutamate receptor, involved in coincidence detection. This may explain the need for distinct types of astrocytes in glycine-rich and glycine-poor regions of the brain.

## Loss of patterning in the oligodendrocyte lineage, and convergence to a single brain-wide intermediate state

Oligodendrocytes wrap myelin sheets around axons to support long-range neurotransmission. We previously described using scRNA-seq (Marques et al., 2016) how the mature cell types are generated from oligodendrocyte progenitor cells (OPCs) that differentiate through a series of intermediate stages including committed oligodendrocyte progenitors (COPs), newly-formed and myelin-forming oligodendrocytes (NFOLs and MFOLs) to acquire one of several myelinating oligodendrocyte fates (MOLs). OPCs remain present in the adult brain, to regenerate myelin as needed. In our previous study, we examined approximately five thousand cells of the oligodendrocyte lineage.

Here, we observed more than two hundred thousand oligodendrocyte lineage cells, yet the overall picture was very similar. The greatly increased sampling depth did not reveal any additional, clearly distinct subtypes of mature oligodendrocytes beyond those we had already described. Furthermore, OPCs remained a single cluster, with only a distinction between cycling and non-cycling OPCs (Fig. 4A-B). In tSNE plots (Fig 4A), a gap still remained between OPCs and COPs, suggesting a rapid transition between those two states (although, tSNE can tend to exaggerate discontinuities). To confirm the relatedness of these two cell types, we examined pairs of mutual nearest neighbors that spanned cluster boundaries; as expected, all links from OPCs extended to COPs, and none to NFOLs (Fig 4A). Cells from OPC, COP and NFOL were intermingled in all tissues, demonstrating a lack of region-specific types of these cells. The exception was COP and NFOL from medulla and pons, which segregated somewhat from other tissues (Fig 4B). However, for technical reasons, those tissues had been analyzed using a different version of the Chromium reagent kit, and thus likely represent a batch effect rather than a genuine biological difference.

**Figure 4.**
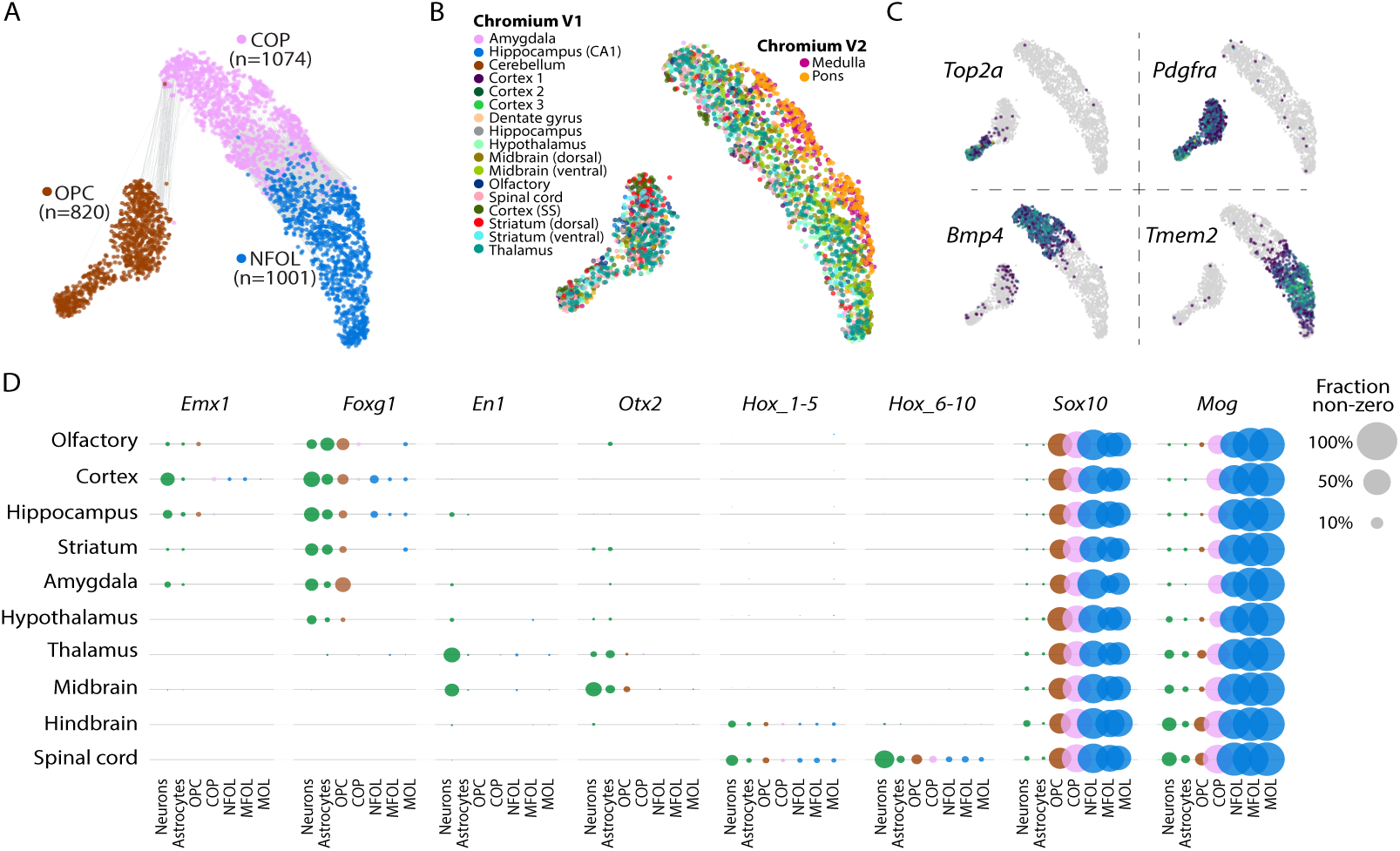
Convergence to a common state at the early stages of oligodendrocytes lineage. (A-B) gt-SNE embedding of the three first stages of the oligodendrocytes lineage OPC, COP and NFOL colored by cluster identity (A) and tissue of origin (B). Edges in (A) connect nodes between mutual neighbors (k = 150), but only if they are from different clusters. (C) Gene expression of selected markers overlaid on gt-SNE embedding. (D) Patterning transcription factor analysis. Circles represent fraction of positive cells in each cluster and brain region.

Thus, regardless of the tissue sampled, OPC, COP and NFOL presented as single, brain-wide common cell types. Since OPCs are the progenitors of the entire oligodendrocyte lineage, this observation demonstrates that the diversity observed among mature oligodendrocytes (Table S3) must be the result of a secondary diversification, not developmental patterning. Oligodendrocyte morphology varies according to the type of axon they myelinate, but transplantation experiments indicate that those differences are plastic (Richardson et al., 2006). This may also explain the graded, interspersed pattern of diversity among mature oligodendrocytes, in contrast to the division into cell types with clear boundaries (molecularly and anatomically) that we observed among astrocytes and neurons. Of mature oligodendrocytes, the spinal cord-enriched MOL3, expressing the serine protease *Klk6* was the most distinct. *Klk6* was recently implicated in the pathogenesis of experimental autoimmune encephalomyelitis, a model of multiple sclerosis in mice (Bando et al., 2018).

OPCs are generated along the length of the neural tube from precursors that are patterned along the antero-posterior axis. This has been demonstrated clearly e.g. by genetic lineage tracing of *Emx1*-positive neural progenitors, which selectively labels forebrain oligodendrocytes (Kessaris et al., 2006). Thus, at some point, cells that later become OPCs must have been molecularly distinct along the antero-posterior axis, for example expressing *Emx1* in the forebrain, *En1* in the midbrain, and *Hox* genes in the hindbrain and spinal cord. Yet, this did not translate into distinct OPC types along the same axis. Clearly, at some point antero-posterior patterning must be lost in the oligodendrocyte lineage. We therefore asked if, despite the lack of clearly distinct subtypes, OPCs, COPs or NFOLs sampled from different tissues retained any traces of patterning gene expression (Fig 4D). We confirmed induction of key transcription factor *Sox10* in OPCs, and of the myelin oligodendrocyte glycoprotein *Mog* in COPs. In the spinal cord, we detected a clear expression of *Hox* genes 6-10 (that is, *Hoxa6*, *Hoxb6*,…,*Hoxd10*), which are responsible for patterning the thoracic spinal cord. These genes were expressed in spinal cord oligodendrocytes only, at levels similar to those observed in neurons and astrocytes, and they remained expressed throughout the oligodendrocyte lineage. Similarly, we found that *Hox* genes 1-5 (that is, *Hoxa1*, *Hoxb1*,…,*Hoxc5*) were expressed in the hindbrain and spinal cord oligodendrocytes, remained expressed into mature oligodendrocytes, and were expressed at levels similar to those in neurons and astrocytes. We conclude that hindbrain and spinal cord oligodendrocytes do retain patterning signals, although this did not translate into clearly distinct hindbrain or spinal cord OPCs, COPs or NFOLs.

In contrast, several transcription factors responsible for patterning the forebrain and midbrain were detected only at very low levels (<1% of cells), if at all, and did not appear in mature oligodendrocytes. At such low levels of expression, it is difficult to rule out a contamination from adjacent neurons, and thus it is possible that they were not expressed at all in the oligodendrocyte lineage. One exception was the forebrain transcription factor *Foxg1*, which was detected in OPCs at levels comparable to neurons and astrocytes. However, its expression was reduced or absent in mature oligodendrocytes.

We conclude that OPCs (and, to a lesser extent, COP and NFOL) may retain a memory of their antero-posterior position, in the form of expression of region-specific transcription factors. However, this does not translate into clearly distinct region-specific cell types, and the memory fades as the cells mature. This is akin to an endogenous reprogramming, analogous to in vitro reprogramming by transcription factors, and shows that cellular states can diverge and then converge. Similar phenomena were recently reported in embryonic stem cells in vitro (Briggs et al., 2017) and in the developing *Drosophila* brain (Li et al., 2017). In the oligodendrocyte lineage, the COP likely represents the convergent cellular state after developmental patterning has been largely lost, but before secondary diversification in response to extrinsic signals has occurred.

## Vascular cells, and a family of broadly distributed mesothelial fibroblasts

A recent paper characterized vascular cells across the murine brain (Vanlandewijck et al., 2018), describing twelve vascular cell types. Our findings agree with these published data, with a few key differences. Like Vanlandewijck et al., we observed distinct endothelial cell types carrying known arterial (e.g. *Bmx*; VECA) and venous (*Slc38a5*; VECV) markers, as well as capillary endothelial cells (VECC) expressing *Meox1*. We found three types of pericytes instead of one (but note that pericytes are notoriously difficult to dissociate from endothelial cells and these subtypes may represent potential endothelial contamination) and a single arterial vascular smooth muscle type (*Acta2*, *Tagln*; VSMCA). Based on the proportion of all cells that were vascular, the midand hindbrain and spinal cord were the most vascularized (Fig. 5B).

**Figure 5.**
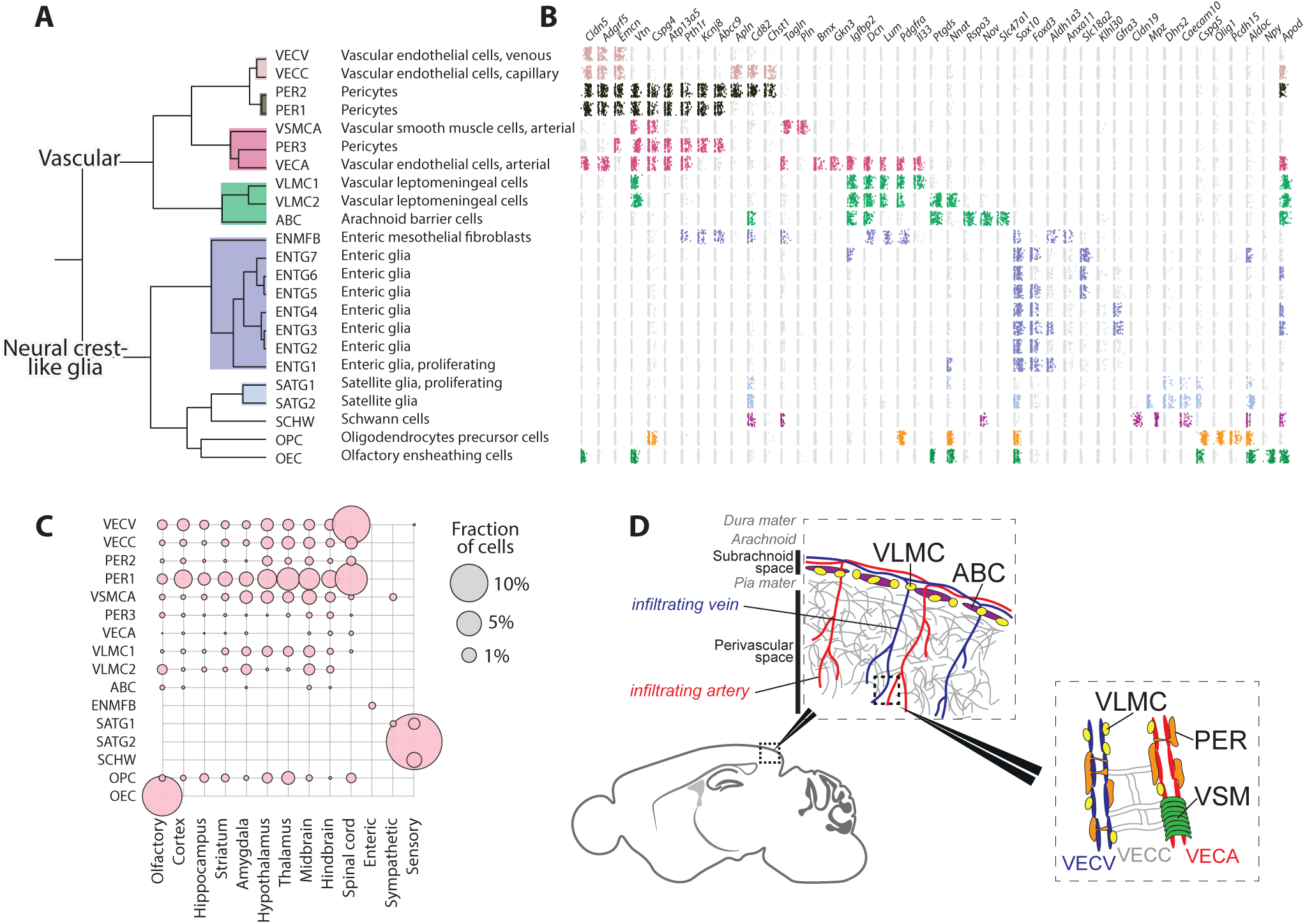
Diversity of the vasculature and neural crest-like glia. (A) Subtree describing the vasculature and neural crest-like glia. (B) Expression dot-plots for marker genes, on log scale and jittered vertically in a uniform interval. Dots are colored only if the trinarization score is positive (posterior probability greater than 0.95), and colors represent the taxonomy rank 4 taxa. (C) The tissue contribution to each cluster, represented by the circle size (enteric glia not shown). (D) Schematic illustration of the approximate position of vascular cell types and the meninges.

Vanlandewijck et al. described two brain fibroblast-like cell types expressing fibril-forming collagens (e.g. *Col1a1*, *Col1a2*), collagen fiber crosslinking proteins (*Lum*, *Dcn*) as well as the platelet-derived growth factor receptor alpha, *Pdgfra*, which were interposed between astrocyte endfeet and vascular endothelial cells. Brain fibroblast-like cells are likely identical to the vascular leptomeningeal cells (VLMCs) that we previously described in mouse CNS (Marques et al., 2016). In the present dataset, we observed four subtypes sharing the canonical markers. Two types were distinguished by expression of genes including the pro-inflammatory cytokine *Il33* (VLMC1) and the Prostaglandin D2 synthetase *Ptgds* (VLMC2), the latter previously shown as the most enriched gene in mouse leptomeninges (Yasuda et al., 2013) (Fig. S6).

Furthermore, we discovered two additional related cell types which shared expression of the canonical VLMC markers. We identified one, ABC, as arachnoid barrier cells, based on the expression of *Abcg2* and *Pgp*. These two genes encode drug and xenobiotic transporters known as BCGP and P-gp, respectively, which are expressed on barrier cells of the arachnoid mater of the meninges (Yasuda et al., 2013). The most specific gene expressed on ABCs was *Slc47a1*, which encodes the multidrug and toxin extrusion protein MATE1, reinforcing the putative function of ABCs to cleanse the cerebrospinal fluid of toxic substances. In contrast to all other VLMC-like cell types, ABCs did not express *Lum*, and showed only very low levels of *Pdgfra*.

The fourth VLMC-like cell type (enteric mesothlial fibroblasts; ENMFB), expressed all the VLMC marker genes, but was found exclusively in the enteric nervous system. This demonstrates that VLMC-like cells are present throughout the body and are not brain-specific. Like the brain, organs of the abdomen are wrapped in protective layers of cells, called the *tunica serosa* and the *tunica adventitia*. These membranes serve protective, lubricating as well as active signalling functions, especially during development. Both serosa and adventitia are made up of mesothelial fibroblasts, but with different properties adapted to freely moving versus rigid organs.

Our observations thus support the view that VLMC-like cells are a family of functionally related (but organ-specific) mesothelial fibroblasts that form protective membranes around internal organs, including the pia and arachnoid membranes of the brain. This is similar to macrophages, which share a common origin in hematopoiesis, but assume organ-specific identities such as the perivascular macrophages and microglia in the brain. The developmental origin of mesothelium outside the brain and heart is unknown (Winters et al., 2012). However, the fact that ENMFBs were obtained by sorting *Wnt1-Cre;R26Tomato* cells indicates that these cells are derived from the neural crest, as has previously been shown for both pia and arachnoid.

## A common regulatory state shared by neural crest-derived glia and oligodendrocyte progenitors

A subtree of the dendrogram in Figure 1 comprised peripheral glia—seven types of enteric glia (ENTG1-7), proliferating (SATG1) and non-proliferating (SATG2) satellite cells of the sensory and sympathetic nervous system and Schwann cells (SCHW)—along with olfactory ensheathing cells (OEC) and oligodendrocyte progenitor cells (OPC) of the CNS.

The function of peripheral glia has been poorly studied, with the exception of Schwann cells, which are the myelinating cells of peripheral nerves. Satellite glia cover the surfaces of sensory and sympathetic neurons and are thought to support their function, but in unknown ways. Satellite glia were enriched in transporters of amino acids (*Slc7a2*), purine nucleobases (*Slc43a3*) and long-chain fatty acids (*Slc27a1*), indicating a role in supporting the metabolism of neurons.

The diversity and function of enteric glia is not known in detail. Enteric glia were very abundant in our dataset (91% of all enteric cells), and almost as diverse as enteric neurons, with seven distinct types. One type (ENTG1) was proliferating (expressing *Top2a*) and could represent a progenitor type. Intriguingly, some enteric glia expressed the vesicular monoamine transporter *Slc18a2* (Fig. S6A), which otherwise loads monoamine neurotransmitters into synaptic vesicles in neurons.

Olfactory ensheathing cells are neural crest-derived (Barraud et al., 2010) cells that ensheath axons of the olfactory sensory neurons, but do not form myelin. Molecularly, they showed a peculiar combination of markers otherwise archetypical of oligodendrocytes (*Plp1*, *Sox10*), pericytes (*Vtn*), endothelial cells (*Cldn5*), neurons (*Npy*) and astrocytes (*Aldoc*), as shown in Figure S6A.

In contrast to all other cell types of this taxon, which are neural crest-derived, OPCs are derived from the neural tube and assumed to be produced by the same progenitors as astrocytes and neurons. Interestingly, however, OPCs share many features of neural crest cells: they require the expression of the two key transcription factors that specify neural crest (*Sox10* and *Sox9*) (Takada et al., 2010), they are highly migratory, and they do not respect developmental borders in the brain. These observations, and the finding that OPCs align molecularly with all the neural crest-derived glia, suggest that they are a neural crest-like type of glia and supports the view that they have a common evolutionary origin with Schwann cells (Kastriti and Adameyko, 2017). Although they are not derived from the physical neural crest, they appear to use similar regulatory mechanisms as neural crest-derived cells. We therefore named this taxon “Neural crest-like glia”.

## Peripheral nervous system

Neurons of the peripheral nervous system segregated molecularly from the CNS, and formed distinct sensory, sympathetic and enteric subdivisions (Fig. 1B and C). Within the peripheral sensory neurons (of the dorsal root ganglia), cell-types were divided into three main branches: peptidergic (8 types), non-peptidergic (6 types) and neurofilament (3 types), which suggests refinements to previous classifications (Li et al., 2016; Usoskin et al., 2014) (Table S2, Fig. 6 and S7).

**Figure 6.**
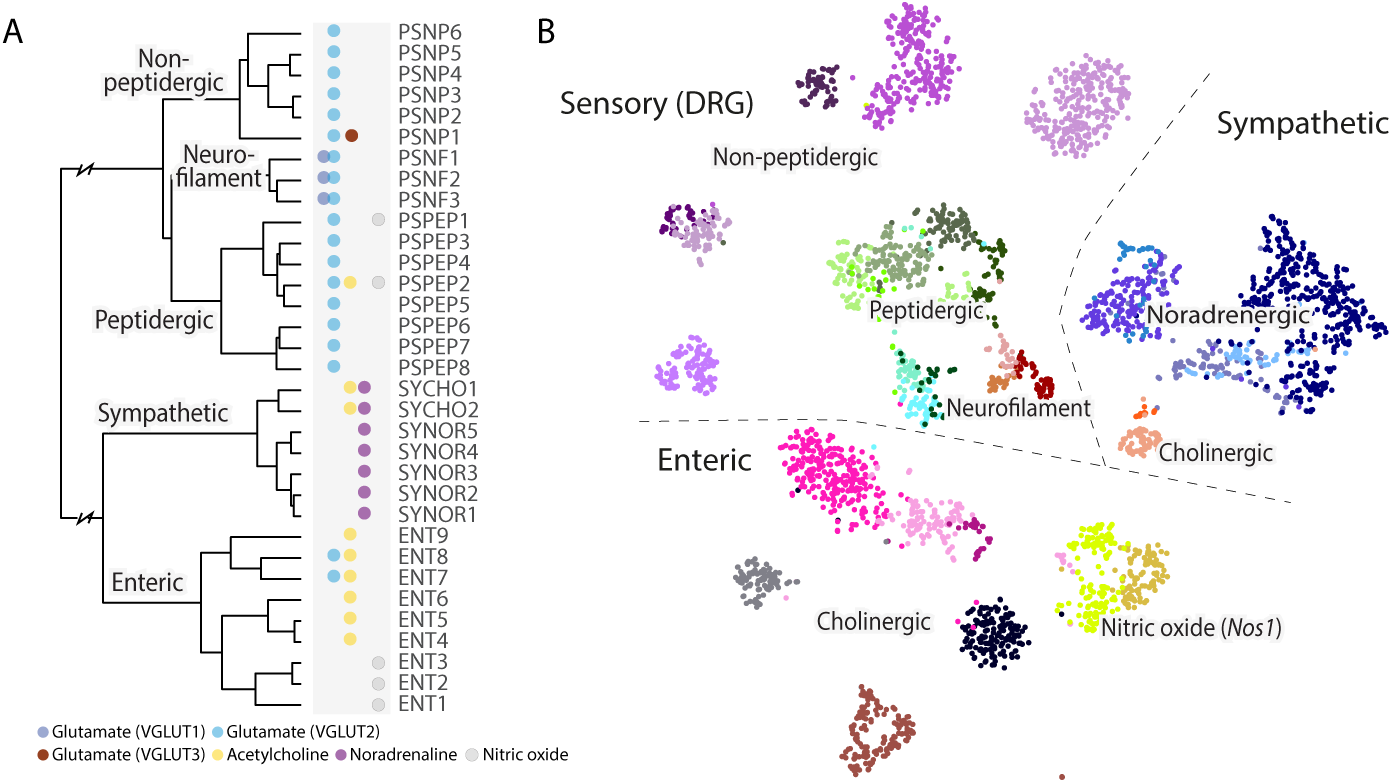
Neurons of the peripheral nervous system. (A) Hierarchical structure of the peripheral nervous system neuronal cell types. Neurotransmitters used by each cell types are indicated by the colored dots next to each leaf. (B) gt-SNE embedding of all related cells demonstrate the diversity and abundance of the different clusters.

Within the sympathetic ganglia we found two cholinergic and five noradrenergic cell types, in agreement with our previous classification (Furlan et al., 2016) (Table S2, Fig. 6 and S7).

The enteric nervous system has not been previously studied in molecular detail using single-cell methods. Here we report on the composition of the myenteric plexus of the small intestine, whereas we did not include cells from the submucosal layer or other regions of the gastrointestinal tract. Based on marker gene expression, morphology, location and projection targets, approximately ten cell types have been previously described (Furness et al., 2014; Qu et al., 2008) in the myenteric plexus of the mouse.

We found nine molecularly distinct neuron types. Although enteric neurons are commonly divided into nitrergic and Calretinin-expressing subtypes, our data indicates that the more natural split is between nitrergic (i.e. expressing the neuronal nitric oxide synthase *Nos1*, ENT1-3) and cholinergic (i.e. expressing *Chat* and *Slc5a7*, ENT4-9) neurons. In addition, ENT7 and ENT8 co-expressed *Slc17a6* (VGLUT2). Many enteric neurons also expressed a variety of neuropeptides (Fig. S7), including *Gal*, *Cartpt*, *Nmu*, *Vip*, *Cck*, and *Tac1*. A more detailed analysis of these neurons will be published elsewhere (manuscript in preparation).

## CNS neurons

A set of 32 clusters formed a subtree of the dendrogram, and included pyramidal cells of the cortex and hippocampus as well as medium spiny neurons of the striatum. We named them *Telencephalon projecting neurons* (expressing high *Ptk2b*, *Ddn*, *Icam5*). The cerebral cortex was the most diverse, with 20 projection cell types, which were glutamatergic (all VGLUT1, but some additionally VGLUT2 or VGLUT3). Closely related were the hippocampal pyramidal cells (three types) and the dentate gyrus granule cells, as well as a single cluster from the basolateral amygdala, all of which— like the isocortex—develop from the pallium. We will discuss the spatial distribution of these cell types in detail below.

The GABAergic medium spiny neurons (MSNs) of the striatum are classified as D1 or D2 type according to the dopamine receptor they express. A long-standing question concerns the diversity of these cell types, in particular relative to structural and functional features of the striatum. For example, dorsal MSNs initiate and control movements, whereas ventral MSNs are involved in motivation, reward, aversion and similar behaviours. We found two D1-type MSNs (MSN1 and MSN4), one enriched in dorsal and one in ventral striatum, as well as two D2-type MSNs (MSN2 and MSN3), also dorsal and ventral, demonstrating a molecular distinction corresponding to the distinct circuits and functions of dorsal and ventral MSNs. In addition, we found putative patch-specific D1/D2-type neurons (expressing *Tshz1*) and matrix-specific D2 neurons (expressing *Gng2*), consistent with staining patterns of these genes in the Allen Mouse Brain Atlas.

Telencephalic inhibitory interneurons, including cells from the olfactory bulb, cortex, hippocampus and thalamus formed a taxon, with the olfactory cells as a separate group. The thalamic inhibitory neurons expressed *Meis2* and shared key transcription factors (e.g. *Dlx1*, *Dlx2*, *Dlx5*, *Dlx6*) with the other cell types in this taxon, as well as with striatum medium spiny neurons, suggests a common developmental origin in the ganglionic eminences.

Most olfactory neurons were GABAergic, consistent with previous work (Nagayama et al., 2014), and two were also dopaminergic. We found no mature glutamatergic neurons in the olfactory bulb. However, two neuroblast types (OBNBL1 and OBNBL2), putatively located in the mitral cell layer, may represent immature versions of the olfactory projection neurons, the mitral and tufted cells. One of them, OBNBL1, expressed the identifying marker of mitral cells, the T-box transcription factor *Tbx21*.

A single taxon collected nearly all cholinergic, monoaminergic neurons (which we identified based on expression of the necessary biosynthesis enzymes and vesicular and reuptake transporters; Fig. S9C) from the whole brain, as well as the peptidergic neurons mainly of the hypothalamus. These included the cholinergic afferent nuclei of cranial nerves III-V (HBCHO4) and VI-XII (HBCHO3), the adrenergic nucleus of the solitary tract (HBADR), the noradrenergic cell groups of the medulla (HBNOR), five serotonergic hindbrain types, two dopaminergic neuron types of the ventral midbrain, and 15 types of peptidergic neurons including those secreting neurotensin (*Nts*), vasopressin (*Avp*), oxytocin (*Oxt*), gonadotropin-releasing hormone (*Gnrh*), galanin (*Gal*), enkephalin (*Penk*), orexin (*Hcrt*), CART peptides (*Cartpt*), thyrotropin (*Trh*), pro-opiomelanocortin (*Pomc*), agouti-related peptide (*Agrp*) and neuromedin (*Nmu*) (Lam et al., 2017; Romanov et al., 2016). Most of these peptidergic cell types were located in hypothalamus, but some were from telencephalon (bed nuclei of stria terminalis and septal nucleus), midbrain (Darkschewitz nucleus) and spinal cord (central canal neurons, see below). The fact that the majority of cholinergic, monoaminergic and peptidergic neurons clustered together suggests a common underlying regulatory state, distinct from that in neurons using canonical neurotransmitters. On the other hand, they still retained their CNS neuron character, and did not intermingle with cholinergic or monoaminergic neurons of the PNS.

We further found 38 excitatory and inhibitory cell types of the diencephalon (thalamus and hypothalamus) and midbrain, forming a unified taxon. These types segregated near-perfectly into glutamatergic (mostly VGLUT1) and GABAergic subsets, but included two cholinergic types (of the red nucleus and the habenula). The thalamus proper contained only glutamatergic neurons, except for the *Meis2*-expressing neurons of the reticular nucleus that forms a capsule around the thalamus. In the midbrain, the superior and inferior colliculi were the most diverse, comprising 17 excitatory (exclusively VGLUT1) and inhibitory (GABA) cell types and spatially distinct distribution. In the ventral midbrain, we identified two types of dopaminergic neurons, one cholinergic and four GABAergic.

In the hindbrain (15 types not including cerebellum), all inhibitory neurons were glycinergic (GLYT1 or GLYT2, or both) and excitatory neurons were a mix of VGLUT1 and VGLUT2. We identified six cell types in the cerebellum, of which five are previously known: Purkinje cells, granular cells, granular layer interneurons, molecular layer interneurons and granular cell neuroblasts. A sixth cell type (MEINH1), curiously, was found in the midbrain but molecularly indistinguishable from cerebellum molecular layer interneurons (CBINH1). It is the only example of a neuronal cell type found in two different and distant regions.

Finally, in the spinal cord we identified 22 cell types, again split into inhibitory (GABAergic or glycinergic) and glutamatergic (VGLUT2) in good agreement with an independent experiment focused on the dorsal horn (Häring et al. in press) (Table S2). In addition, here we identified central canal neurons (SCINH11), known as cerebrospinal fluid-contacting neurons, which expressed transcription factors *Gata2* and *Gata3* (Fig. S10) (Petracca et al., 2016). They also specifically expressed polycystin-like genes (*Pkd1l2* and *Pkd2l1*) which encode a mechanosensory protein complex that detects fluid flow, and *Espn*, which encodes an actin bundling protein with a major role mediating sensory transduction in mechanosensory cells. Thus, central canal neurons are likely specialized cells that monitor cerebrospinal fluid flow.

## Spatial distributions reflect molecular diversity

Given the importance of location for neuronal function, we wanted to assign a spatial distribution to each cell type. The Allen Mouse Brain Atlas provides systematic high-quality information about gene expression, based on in situ hybridization. The data is available both as images and in the form of three-dimensional volumetric maps. We computed the spatial extent of each cell type by correlating volumetric and RNA-seq gene expression, using only cell type-specific genes as determined by a significant enrichment score. That is, using only cell type-enriched genes, for each voxel we computed the correlation of RNA-seq expression values with in situ hybridization expression values. The resulting data was visualized as three-dimensional density maps, expected to peak in regions where each cell type was abundant. We used the Allen Mouse Brain Atlas anatomical ontology to name the top-ranked anatomical regions for each cell type.

Note that this approach will clearly fail in certain cases. First, if the cell type is not located within the CNS, then the resulting density map will be meaningless, and we therefore do not provide maps for peripheral neurons. Second, many enriched genes can be shared between a set of related cell types (e.g. cortical pyramidal cells), with only a smaller number of highly specific genes; in those cases, we expect the density map to show both the overall distribution of related cell types, as well as a stronger peak at the location of the specific cell type. To alleviate this concern, we carefully curated the location of each cell type by manual inspection of individual in situ hybridization images (available in Table S3, column “Probable location”). We sought to provide the most plausible location for each cell type, rather than refraining from assigning a location when it was uncertain.

Inspecting the resulting cell type distribution maps, we found reassuringly that the automatically assigned locations corresponded well with the known source of the cells. For example, cortical and hippocampal projection neurons were assigned to cortex and hippocampus as expected (Fig. S8). But the spatial maps provided much more detail: for example, the distinction between CA1 and CA3 pyramidal cells was clear (Fig. S8, right), and cortical pyramidal cells could be assigned highly specific distributions across the cortical surface (Fig. S8, left) and layers. Interestingly, the spatial distribution of cortical pyramidal neurons correlated with their molecular similarity. For example, pyramidal neurons of the piriform and entorhinal cortex, as well as the subiculum, were molecularly closely related (shown by their forming a separate subtree of the dendrogram) as well as spatially aligned. Similarly, the pyramidal cells of the neocortex were arranged by molecular similarity in layer order (i.e. layers 2/3, layer 4, layer 5, layers 6/6b). Notably, this also corresponds to their order of development during embryogenesis.

Beyond the cortex, many cell types were assigned to very specific locations, greatly aiding interpretation of the data. For example, midbrain dopaminergic neurons (MBDOP2) were found in the substantia nigra and ventral tegmental area (Fig. 7). One of the most complex regions was the superior and inferior colliculi, to which 17 celltypes were assigned (ten excitatory, seven inhibitory). Spatial distribution maps are provided for all CNS neurons at the companion wiki web site (Box 1).

**Figure 7.**
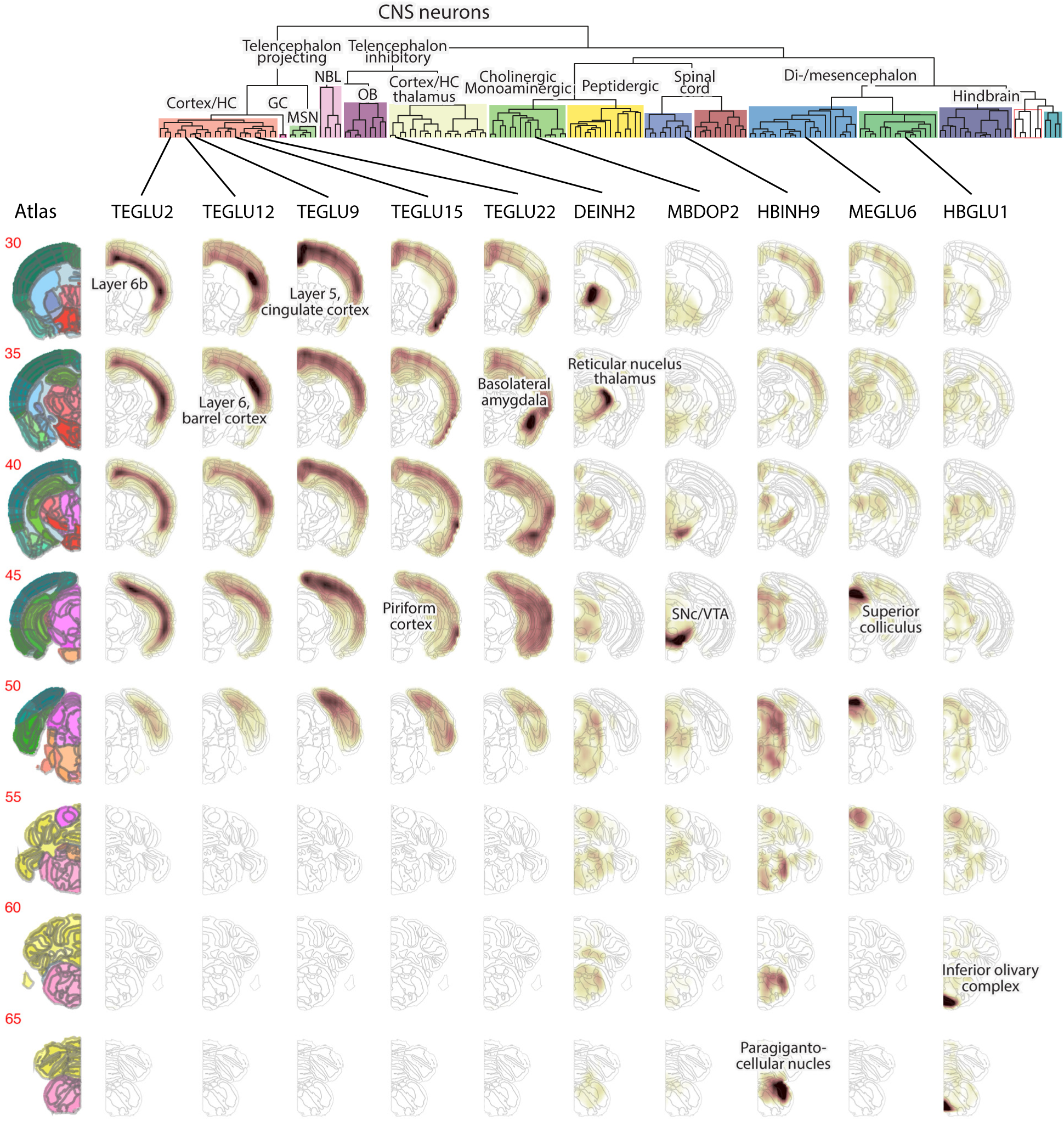
Neuronal cell types are spatially restricted. Examples of inferred spatial distributions for cell types across the brain. Left column show reference images from the Allen Brain atlas. Each row shows one coronal section, ordered rostro-caudally, and each column shows one cluster as indicated at the top. For every cluster and every voxel, the correlation coefficient is depicted by the colormap (dark high, white low). Labels indicate the top-scoring anatomical unit for each cluster.

## Drivers of neuronal and glial diversity

In order to better understand the forces that drive gene expression diversity in the mammalian nervous system, we next examined the expression of neurotransmitters and neuropeptides. How do diverse neurotransmitters cooperate or interact with each other? And more generally, how do neuropeptides overlap with classical neurotransmitters and with each other, in different contexts across the nervous system? We examined the co-expression of neurotransmitters, while retaining information about the tissue compartment (Figure S9A). While VGLUT2 was the only neurotransmitter expressed in all compartments, GABA contributed the larger number of cell types and was mostly concentrated in the forebrain. GABAergic and glutamatergic (VGLUT1 and VGLUT2) neurotransmission was mutually exclusive; we did not find a single cell type anywhere in the nervous system that expressed both. Glutamatergic neurons in the telencephalon all used VGLUT1, with some additionally using VGLUT2, whereas in more caudal regions, VGLUT2 dominated. Interestingly, the boundary that separated VGLUT1 dominance from VGLUT2 dominance appeared to be the telencephalon-diencephalon border, analogous to the separation of the two major types of astrocytes at this same boundary (although, both were expressed in the thalamus).

In contrast, the atypical vesicular glutamate transporter VGLUT3 was often co-expressed (Figs. 1C and S9A) with cholinergic and monoaminergic neurotransmitters (12 cell types) and more rarely alone (1 cell type) or with the other VGLUTs or GABA. This supports the notion that VGLUT3 plays a distinct role in cell types that release non-canonical neurotransmitters.

Acetylcholine occurred alone, or together with noradrenaline (in the sympathetic nervous system) or glutamate, but never with GABA or glycine. Similarly, noradrenaline co-localized with acetylcholine, whereas serotonin occurred only alone or with VGLUT3. In contrast, the gaseous neurotransmitter nitrix oxide (i.e. *Nos1*) was detected throughout the nervous system, and did not combine preferentially with (or avoid) any other neurotransmitter (Figs. 1C and S9A).

Examining the co-expression matrix of individual genes encoding neurotransmitter enzymes, vesicular and reuptake transporters, and neuropeptides, we found stereotyped combinatorial patterns assigned to specific compartments of the nervous system (Figure S9C). This analysis demonstrates how the rules governing gene co-expression can vary between brain regions. For example, somatostatin (*Sst*) is a canonical marker of inhibitory neuronal subtypes in the forebrain, but was widely expressed in excitatory neurons in the spinal cord, hindbrain and di-mesencephalon. Moreover, *Sst* was also expressed in combination with *Fev* (serotonin, hindbrain), *Dbh* (noradrenalin, PNS) or the neuropeptide *Trh* (hypothalamus). *Pvalb*—another canonical marker of forebrain inhibitory cells—was also expressed in excitatory neurons in the mid - and hindbrain. These results demonstrate that neuropeptides, neurotransmitters, calcium-binding proteins, and other neuronal molecules are used in a highly modular fashion and serve different functions in different contexts.

Expanding the analysis to all genes, we note that the dendrogram and taxonomy (Fig. 1C, S2 and S3) reflect systematic patterns of shared and unique gene expression. However, while some subtrees and taxa reflect biologically natural categories, with shared molecular properties, others may be more diverse and lack such shared features. For example, the distinction between CNS and PNS may reflect shared properties primarily among PNS neurons, primarily among CNS neurons, or both. To gain more insight into these patterns, we systematically searched for genes that were expressed ubiquitously in one set of cell types, but absent from most other cell types (Fig. S9B, Table S4). We used stringent statistical criteria (Fisher’s exact test with 5% false discovery rate using Benjamini Hochberg correction), and retained only genes that were detected in more than 70% of cell types within the set, but less than 10% of clusters outside it. We note that, since the scRNA-seq data is far from 100% sensitive, our estimates of class-specific genes are very conservative and more sensitive measurements may uncover additional such genes.

Comparing first neurons to all non-neurons, corresponding to the first split of the dendrogram, we found 205 pan-neuronal genes. These included well-known neuronal markers (e.g. the kinase *Camk2b*, the beta-3 tubulin *Tubb3* and the membrane glycoprotein *Thy1*); transcription factors *Myt1l*, *Ncoa7*, *Mafg* and *Zcchc18*; synaptic proteins (synaptosome proteins *Snap25*, *Snap47* and *Snap91*, synapsins *Syn1* and *Syn2*, synaptogyrin *Syngr1*, synaptojanin *Synj1*, synaptophysin *Syp*, synaptotagmins *Syt1* and *Syt4* and neurexin *Nrxn3*); stathmins (*Stmn2* and *Stmn3*); as well as the RNA-binding proteins *Elavl4* (also known as *HuD*), *Rbfox1*, *Rbfox2* and *Rbfox3* (also known as *NeuN*).

In contrast, examining glia as a group (but excluding microglia), we found only six pan-glial genes, including *S100a1*, showing that macroglia is not a natural category of cells with shared properties. However, there were 74 pan-astrocyte genes (e.g. *Aqp4*, *Sox2* and *Sox9*), 135 pan-oligodendrocyte genes (e.g. *Sox10*) and 51 genes shared across neural crest-like glia (e.g. the transcription factors *Sox10* and *Hes1*, and the myelin protein zero-like protein *Mpzl1*, related to the Schwann-cell specific *Mpz*). Thus each major glial lineage is characterized by distinct gene modules, with very few genes in common across PNS and CNS glia.

We found 111 genes that were ubiquitous in PNS but not CNS neurons. The top hits included the homeobox transcription factor *Tlx2* (also known as *Enx*, *Ncx* and *Hox11l1*), the intermediate filament cytoskeletal protein peripherin (*Prph*) and the phosphoinositide-interacting membrane protein *Pirt*. *Tlx2*-null mice show myenteric neuron hyperplasia and megacolon (Shirasawa et al., 1997), supporting a fundamental role in enteric neuron specification, whereas *Pirt* knockout animals are viable, fertile and show normal appearance and behaviour (Kim et al., 2008).

In contrast, we found only ten genes that were expressed in most CNS but not PNS neurons, including the NMDA glutamate receptor subunit *Grin2b*, the potassium/choline transporter *Slc12a5*, the tyrosine kinase *Matk*, and the fractalkine ligand *Cx3cl1* (which binds to the CX3C chemokine receptor, encoded by *Cx3cr1*, found on immune cells in the brain). Thus, while the PNS is a natural biological category that shares expression of many genes, the CNS is too diverse to be considered as a unit.

FOCUSING NEXT ON CNS NEURONS, we searched for the genes that contributed the most to CNS neuronal diversity. We collected the top ten most highly enriched genes in each cell type, reflecting both high expression and high specificity (in contrast to the pan-CNS genes analyzed above). A gene set enrichment analysis against the Gene Ontology (Fig. 8A) (Huang et al., 2009) pointed to four clear categories of genes: those that establish cell identity (e,g, transcription factors, developmental genes), membrane conductance (e.g. ion channels, calcium binding proteins), neurotransmission (e.g. neurotransmitter synthesis enzymes, transporters, neuropeptides and their receptors), and synaptic connectivity (e.g. synaptic and cell junction proteins). These findings point to the specific functions that differ between neuronal types (connectivity, electrophysiology and neurotransmission), and to the underlying regulatory machinery (transcription factors).

**Figure 8.**
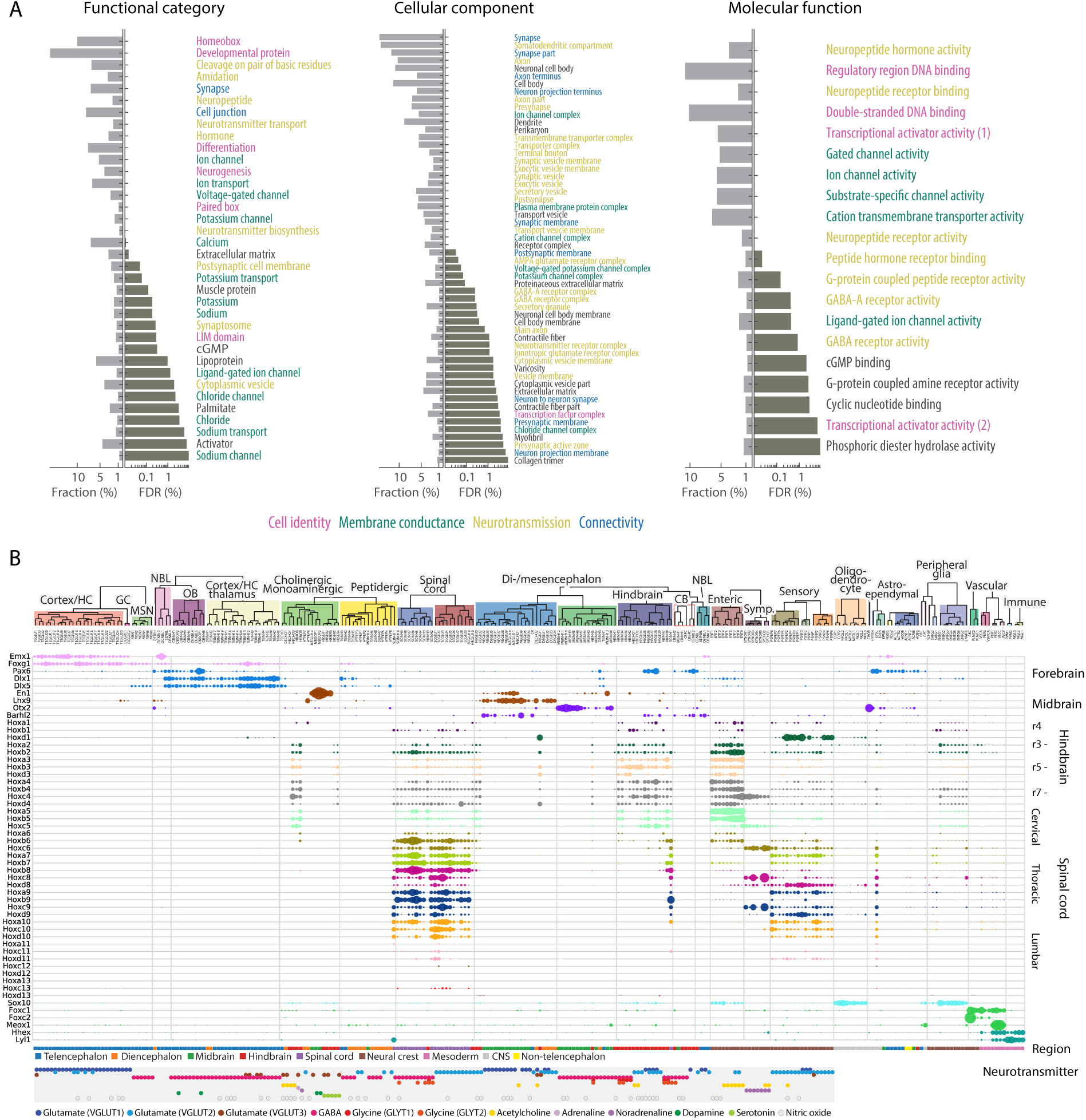
Drivers of cellular diversity. A) Gene ontology analysis of the most highly enriched genes in CNS neuronal clusters. Each panel shows the significantly (false discovery rate, FDR < 10%) enriched terms, ranked by FDR. Bars show the percentage of all genes (belonging to each term) that were enriched, and the FDR. Colors indicate major categories of terms, as indicated below the figure. (B) Gene expression of developmental patterning transcription factors is shown along the cell types taxonomy. Each row represents one transcription factor and columns represent clusters. Circles represent mean expression values, proportional to area. Genes are sorted according to their expression pattern, with *Hox* genes sorted rostro-caudally. Labels on the right indicate the approximate anatomical extent of the expression of corresponding *Hox* genes.

The gene family that best distinguished CNS neuron classes (defined by the “functional category” label) was homedomain transcription factors (Fig. S10), consistent with an important role in specifying and maintaining neuronal cell types. Other transcription factor families showed less specificity in the CNS, but were more variably expressed in the PNS and among glia and vascular cells.

Many homeodomain transcription factors are involved in dorso-ventral and antero-posterior patterning (as well as the specification of e.g. the neural crest). Although patterning takes place during embryogenesis, we reasoned that significant traces of patterning gene expression might remain, and could explain the observation that the nervous system was molecularly organized according to developmental origin. In agreement with this prediction, we found that *Hox* genes were expressed in cell types derived from the hindbrain, spinal cord and the peripheral nervous system (Fig. 8B). For example, spinal cord cell types expressed *Hoxa1* (rhombomere 1) through *Hoxd10* (thoracic bordering on lumbar), with additional expression of lumbar *Hox* genes in some cell types. Most spinal cord cell types appeared to express a similar range of *Hox* genes, corresponding to cervical and thoracic levels and indicating that they were not segment-specific (rather, the subtypes corresponded to dorso-ventral layers). However, three inhibitory types expressed *Hoxa11*, *Hoxb11* and *Hoxc11*, indicating a more lumbar extent. Two cell types located in the medulla (HBINH9 and HBGLU10) intermingled with spinal cord cell types in the dendrogram, but did not express *Hox* genes beyond *Hoxb7*. These cell types were presumably derived from the most posterior part of the medulla, explaining their spinal cord-like character, yet they retained proper medulla patterning gene expression.

Enteric neurons and glia of the small intestine, which are believed to develop from the vagal (neck) neural crest, both expressed a *Hox* code broadly consistent with a vagal origin. However, we also noticed some expression of more thoracic Hox genes, indicating a broader origin of these cells.

Sensory and sympathetic neurons, as well as satellite glia, expressed *Hox* genes from all rostrocaudal levels (lumbar cells were not analyzed in the sympathetic nervous system). However, curiously, sympathetic neurons showed highly preferential expression from the *HoxC* cluster only. This is reminiscent of the role of the *HoxD* cluster during digit formation (Deschamps, 2008), and suggests that the *HoxC* cluster may be involved in the specification of distinct sympathetic cell types along some spatial axis.

*Hox* genes are not expressed in the forebrain and midbrain. Nevertheless, as in the hindbrain and spinal cord, forebrain cell types retained patterning gene expression. For example, the forebrain patterning gene *Foxg1* was found in all forebrain neurons, as well as in telencephalon-specific astrocytes. Dorso-ventral patterning was also preserved: the dorsal gene *Emx1* was expressed in cortical, hippocampal and striatal projection neurons, whereas ventral *Dlx1* and *Dlx5* were found mainly in inhibitory neurons of the same tissues.

Among cholinergic and monaminergic neurons, which did not align molecularly with their tissue of origin (Fig. 1), those sampled from the medulla retained patterning gene expression (e.g. *Hox* genes in HBSER4, see Table S3 and Fig. 8) whereas those from the pons did not (e.g. HBCHO4, afferent nuclei of cranial nerves III-V).

## Discussion

We have described the molecular architecture of the mammalian nervous system, based on a systematic survey using single-cell RNA sequencing.

Although we present a comprehensive analysis, our data has several limitations. First, there were technical and experimental limitations as detailed above, including doublets, sex-specific gene expression and low-quality cells. Second, we sampled only a little more than half a million cells across the nervous system, and deeper sampling is likely to reveal additional structure that was obscured in the present study. Similarly, we used relatively shallow sequencing, and deeper sequencing using more sensitive RNA-seq methods is likely to resolve more subtypes. Third, some cell types may have been lost to differential survival or size selection biases (for example, Purkinje cells were likely undersampled here due to their size). Fourth, we have performed a very conservative clustering, designed to reveal clearly distinct major cell types, but did not analyze the substantial remaining heterogeneity within clusters. Finally, we have described only molecular cell types, but the task of linking molecular properties to functional, anatomical, morphological and electrophysiological properties remains.

We suggest that the diversity of gene expression patterns in the nervous system can be understood through three major principles.

First, major classes of cells—e.g. neurons, astrocytes, ependymal cells, oligodendrocytes, vascular and immune cells—are distinguished by large sets of class-specific genes that implement the specific function of each class of cells. For example, neurons share an extensive gene program involving synaptic, cytoskeletal and ion channel genes, while oligodendrocytes express gene programs required for generating myelin. Multiple levels of hierarchical subdivision exist within these classes; for example, within neurons, neurotransmitter phenotype showed a modular and highly regulated pattern of expression.

Second, some—but only some—cell classes show area-specific patterns of gene expression that likely reflect their developmental history. This trend was strongest amongst neurons, astrocytes, and ependymal cells; by contrast, oligodendrocytes, vascular, and immune cells exhibited similar gene expression patterns across brain regions. The territories defining these gene expression domains corresponded closely to those marked out by embryonic morphogens, and spatial differences in adult expression patterns correlated with persistent expression of developmental transcription factors. This suggests that transcription factor networks induced in early development by local morphogens result in heritable regulatory states which in turn are relayed into the diversification of terminal neuronal and astrocytic types specific to each brain region. The fact that oligodendrocytes did not show similar spatial patterns— despite being derived from the same initially patterned neural tube as neurons and astocytes—suggests there is a prevalent loss of regional patterning in the oligodendrocyte lineage, presumably because region-specific patterning is transient and not converted to permanent states in these lineages.

Third, a secondary diversification, more graded and less region-specific, results from interaction with the local environment, and likely reflects inducible gene regulatory networks that respond in graded and transient fashion to local molecular cues. This was observed most clearly in the oligodendrocyte lineage, but likely occurs to some extent in all lineages.

It remains unclear why the initial patterning is retained by neurons and to some extent astrocytes, but not by oligodendrocytes. Among CNS neurons, we found that four main categories of genes drive neuronal diversity: those involved in cellular identity (transcription factors), connectivity (synaptic proteins, junction proteins), neurotransmission (neurotransmitters, neuropeptides), and membrane conductance (ion channels, calcium-binding proteins, solute carriers). But synaptic connectivity, neurotransmission and membrane conductance are uniquely neuronal properties, and their diversity between regions in consistent with the diverse computational roles of each neuronal circuit. Conversely, the relative homogeneity of oligodendrocytes points to a common function, myelination, across all regions. The intermediate behaviour of astrocytes is therefore consistent with the emerging view that they are not simply support cells, but play an active role in computational processing (Henneberger et al., 2010).

Our atlas can be used to identify genes and gene combinations unique to specific cell types, which in turn can be used to genetically target cells for visualization, ablation, optogenetic manipulation, gene targeting and more. Surprisingly, we found that two genes were sufficient to uniquely target most cell types in the entire nervous system, and none required more than three genes. These findings provide a powerful starting point for precise genetic manipulation of defined cell types in the mouse nervous system.

The atlas will also help us understand the function of specific genes, for example those implicated in disease (Skene and Grant, 2016). This can lead to actionable hypotheses on the mechanism of disease as well as identifying the relevant cell types to generate mouse models of human disease. Similarly, one can use the atlas to find, across the entire nervous system, those cell types likely to respond to a drug (with known target). This will be important to advance our understanding of the specificity of drugs and their potential off-target effects.

In summary, we provide a resource and an initial analysis revealing key principles of the molecular diversity and composition of the mammalian nervous system.

## Acknowledgements

We thank Maayan Harel for the drawing in Fig. 1A, Igor Adameyko, Christer Betsholtz and Michael Vanlandewijck for stimulating discussions, the National Genomics Infrastructure at Science for Life Laboratory for sequencing services, and the Eukaryotic Single-cell Genomics core facility at Science for Life Laboratory for use of equipment. This work was supported by grants from the Knut and Alice Wallenberg Foundation (2015.0041), the Swedish Foundation for Strategic Research (RIF 14-0057 and SB16-0065), and the Wellcome Trust (108726/Z/15/Z) to S.L.

## Methods

### Animals

Table S1 details the animals used per experiment. In summary, male and female mice were postnatal ages P12-30, as well as 6 and 8 weeks old. We mainly used wild type outbred strains CD-1 (Charles River) and Swiss (Janvier). *Wnt1-Cre:R26Tomato* (C57Bl6J background) (Danielian et al., 1998; Madisen et al., 2010) were used to isolate peripheral and enteric nervous system, and *Vgat-Cre:tdTomato* (heterozygous for *Cre* and homozygous for *tdTomato*; mixed CD-1, C57BL/6J background) (Ogiwara et al., 2013) to isolate inhibitory neurons (vesicular GABA transporter, *Slc32a1*). All experimental procedures followed the guidelines and recommendations of Swedish animal protection legislation and were approved by the local ethical committee for experiments on laboratory animals (Stockholms Norra Djurförsöksetiska nämnd, Sweden).

### Single-cell dissociation (brain)

Single cell suspensions of all brain regions, i.e. all regions except spinal cord, sympathetic and enteric nervous system as well as dorsal root ganglia, were prepared as described previously (Hochgerner et al., 2018). Briefly, mice were sacrificed with an overdose of isoflurane, followed by transcardial perfusion with artificial cerebrospinal fluid (aCSF, in mM: 87 NaCl, 2.5 KCl, 1.25 NaH2PO4, 26 NaHCO3, 75 sucrose, 20 glucose, 1 CaCl2, 7 MgSO4). The brain was removed, 300μm vibratome sections collected and the regions of interest microdissected. The pieces were dissociated using the Worthington Papain kit, with 25-35 min enzymatic digestion, as needed, followed by manual trituration using fire polished Pasteur pipettes and filtering through a 30μm aCSF-equilibrated cell strainer (CellTrics, Sysmex). Importantly, aCSF equilibrated in 95% O2 5% CO2 was used in all steps, and cells were kept on ice or at 4°C at all times except for enzymatic digestion.

### Single-cell dissociation (spinal cord, sympathetic and dorsal root ganglia)

CD-1 mice (DRG and spinal cord) or *Wnt1-Cre:R26RTomato* mice (sympathetic) were sacrificed and tissues of interest collected in freshly oxygenated, ice cold aCSF (see above). Sympathetic (SG) and dorsal root ganglia (DRG) were dissected and dissociated as described before (Furlan et al., 2016), with minor modifications. Briefly, following dissection (DRG: ~30 ganglia collected in total from cervical 1 - lumbar 6; SG: thoracic 1 - 12 and stellate), the ganglia got transferred into a 3cm plastic dish with 2.7ml of pre heated (37°C) digestion solution (400µl TrypLE™ Express (Life Technologies), 2000µl Papain (Worthington; 25U/ml in aCSF), 100µl DNAse I (Worthington; 1mM in aCSF) and 200µl Collagenase/Dispase (Roche; 20mg/ml in CS)). Non-ganglia tissue was removed from the ganglia. After 30 min incubation at 37°C, ganglia were triturated with 0.5% BSA-coated glass Pasteur pipette (flamed to 70% of original opening). DRG were also carefully ripped open by using fine forceps to make cells more accessible for the enzymes. This procedure was repeated every 20-30 min using Pasteur pipettes with decreasing diameter appropriate to the dissociation state. Depending on the dissociation progress 50µl of Collagenase/Dispase (20mg/ml) and 100µl of TrypLE solution was added.

Dissociation of the spinal cord followed the procedure described in Häring et al., (2018, in press). In short, following the isolation of grey matter (from cervical to sacral levels), the tissue was transferred into a 3cm plastic dish with 2.5ml of pre heated (37°C) digestion solution (300µl TrypLE™ Express (Life Technologies), 2000µl Papain (Worthington; 25U/ml in aCSF), 100µl DNAse I (Worthington; 1mM in aCSF) and 100µl aCSF. Meninges were removed and the grey matter cut into pieces 1-2mm2. After 30 min incubation at 37°C, pieces were triturated with the first Pasteur pipette (see above). This procedure was repeated every 20min using Pasteur pipettes with decreasing diameter appropriate to the dissociation state. Depending on the progress of spinal cord dissociation, 100µl of TrypLE solution was added.

As soon as all ganglion or spinal cord pieces were dissociated (DRG, SG: ~1.5-2h; Spinal Cord: 45-60min), the cell suspensions were filtered using a 40µm cell strainer (FALCON) and collected in a 15ml plastic tube. The digestion solution was diluted with 3ml aCSF and centrifuged at 100g for 4min at 4°C. The supernatant was removed and the pellet resuspended in 0.5ml aSCF and 0.5ml complete Neurobasal medium (Neurobasal-A supplemented with L-Glutamine, B27 (all Gibco) and Penicillin/Steptamycin (Sigma)). The cell suspension was carefully transferred with a Pasteur pipette and layered on top of an Optiprep gradient: 90µl (DRG) or 80µl (SG) Optiprep Density Solution (Sigma) in 455µl aCSF and 455µl complete Neurobasal; and for spinal cord 170µl of Optiprep in 915µl aCSF and 915µl complete Neurobasal. The gradient was centrifuged at 100g for 10min at 4°C, the supernatant removed until only 100µl remained and 10µl DNaseI added to avoid cell clumping.

### Single-cell dissociation (enteric nervous system)

*Wnt1-Cre;R26RTomato* mice were killed by cervical dislocation followed by dissection of small intestine. During all steps the tissue was kept in aCSF (in mM: 118 NaCl, 4.6 KCl, 1.3 NaH2PO4, 25 NaHCO3, 20 glucose, 7 mM CaCl2 and MgSO4) equilibrated in 95% O2 5% CO2 for 30 min before use and held on ice. The small intestines of male and female (P21) mice were cut in 5cm pieces and flushed clean with ice-cold aCSF using a blunt 20G needle attached to a 20ml syringe. The mesentery was removed, the pieces opened lengthwise along the mesenteric border and pinned with the mucosa side down on a Sylgaard (Dow Corning) covered dissection dish. The outer smooth muscle layers, containing the myenteric plexus were peeled off from the submucosa using forceps. The tissue was digested in 1,5 mg/ml Liberase™ (Grundmann et al., 2015), 0.1 mg/ml DNAseI and 1xAntibiotic-Antimycotic (ThermoFisher) in aCSF at 37°C for 1h, with shaking of the tube every 15 min. The cells were gathered by centrifugation at 356g for 5min followed by incubation in TrypLE for 30 min. The suspension was washed in aCSF, centrifuged at 356g for 5 min and resuspended in aCSF, 1% BSA. After manual trituration using BSA-coated fire-polished Pasteur pipettes with decreasing opening size, the single cell suspension was filtered through 70μm filter (Miltenyi Biotec) and cleaned of debris by centrifugation through 1 ml FBS at 800g for 10min. The cells were resuspended in oxygenated aCSF, 1%BSA and filtered through a 30μm filter (Miltenyi Biotec). Tom+ cells were FAC sorted on a BD FACSAria II and collected in ice cold aCSF.

### Single-cell RNA-seq (10x Genomics Chromium)

The majority of sampling was carried out with 10X Genomics Chromium Single Cell Kit Version 1, although part of the hindbrain sampling was done in Version 2 (Table S1). Suspensions were prepared as described above and diluted in aCSF, to concentrations between 300-1000 cells/μl (listed in Supplementary Table S1), and added to 10x Chromium RT mix to achieve loading target numbers between 2500-8000 (V1 kit) or 7000-10,000 (V2 kit), as indicated. For downstream cDNA synthesis (12-14 PCR cycles), library preparation, and sequencing, we followed the manufacturer’s instructions.

### RNAScope

CD-1 mice (Charles River) were killed with an overdose of isoflurane and transcardially perfused with artificial cerebrospinal fluid. Brains were dissected out, snap frozen in OCT on a bath of isopentane with dry ice and stored at –80°C. Fresh frozen sagittal whole-brain sections (including the olfactory bulb, SVZ, hippocampus and cerebellum) of 10 µm thickness were cryosectioned and stored at –80°C. Sections were thawed just prior to staining and fixed with 4% PFA for 15 min followed by rinsing in PBS. RNAScope in situ hybridizations were performed according to the manufacturer’s instructions, using the RNAScope Multiplex Fluorescent kit (Advanced Cell Diagnostics) for fresh frozen tissue, as previously described(Hochgerner et al., 2018). A 10 min treatment in SDS (4% in 200 mM sodium borate) was added in the protocol after the Protease IV incubation. Following probes with suitable combinations were used (indicated with gene target name for mouse, respective channel and catalogue number from Advanced Cell Diagnostics): Mfge8 408771, Agt 426941-C2, Aqp4 417161-C3, Slc6a9 525151, Slc6a11 492661-C3, Islr 450041 and Gdf10 320269-C2. All sections were mounted with Prolong Diamond Antifade Mountant (P36961, ThermoFisher Scientific). Imaging was carried out on a Nikon Ti-E epifluorescence microscope (Nikon) at 10X magnification.

### Cytograph pipeline

Chromium samples were sequenced, typically one sample per lane, per the manufacturer’s instructions with one 98 bp read located near the 3′ end of the mRNA. Illumina runs were demultiplexed, aligned to the genome and mRNA molecules were counted using the 10X Genomics cellranger pipeline.

Each raw Chromium sample was manually inspected after sequencing. Samples that showed no obvious structure in their t-SNE plots (generated automatically by the Chromium cellranger pipeline) were excluded from further analysis. The complete list of input samples is given in Table S1.

All subsequent analyses were automated in the cytograph library and adolescent-mouse pipeline, freely available as open source. Cytograph evokes both the fact that our cell type clustering and visualizations are graph-based, and the fact that the pipeline itself is organized as a directed acyclic graph.

Our pipeline is based on Luigi (Spotify), a Python-based software that orchestrates a set of tasks with dependencies. Each task takes zero or more input files, and generates exactly one output file. Luigi automatically determines which outputs are missing, and the order in which tasks have to be executed to generate them. It can also allocate independent tasks in parallel, to increase throughput.

### Quality controls

Cells with less than 600 detected molecules (UMIs), or less than 1.2-fold molecule to gene ratio, were marked invalid. Genes detected in fewer than 20 cells or more than 60% of all cells were marked invalid. These filters were applied separately to each input file.

### Preliminary exploratory analysis

In preliminary analyses, we explored a large number of approaches for dimensionality reduction, manifold learning, clustering and differential expression analysis methods, in order to get a deep preliminary understanding of the dataset.

For normalization and noise reduction, we tried simple things like mean-centering, normalization to a common molecule count, standardization (division by the standard deviation) and log transformation; we also explored MAGIC (a method that imputes expression based on neighbors in the KNN graph) and diffusion maps.

For manifold learning, we projected the high-dimensional dataset either to a graph (e.g. of k nearest neighbors KNN, and variants such as mutual nearest neighbors) or to two or three dimensions (using PCA, t-SNE, SFDP). We also combined these approaches, first projecting to a graph, then calulating distances on the graph (e.g. Jaccard distance, or multiscale KNN distance; see below), then using those distances to project to 2D space using graph-t-SNE (gt-SNE; see below).

For clustering, we explored standard methods such as K-means (and iterative K-means) in PCA space, as well as graph-based algorithms (Louvain community detection) and density-based algorithms in 2D or 3D projections (e.g. DBSCAN, HDBSCAN).

The final algorithm choices below reflect what we learned in this exploratory phase.

### Preliminary clustering and classification

We extensively mined clusters obtained in preliminary analyses and found that they largely corresponded to known and putative cell types, broadly consistent with previous data. Some clusters were also clearly derived from doublets, expressing contradictory markers e.g. from neurons and vascular cells.

With any type of clustering the choice of feature space is crucial. For preliminary clustering, we used genes informative across the entire set of cells, projected by PCA. This would be expected to be suitable for finding major cell types, but would not be optimal for finding finer subdivisions among cells of the same kind (e.g. interneurons in a dataset containing both neurons, vascular cells and glia). For example, running Louvain clustering on the full dataset resulted in only 44 clusters, compared to the 265 found by the multi-level, iterative approach described below.

We decided to first split cells by major class. In order to split the data, and to reject many doublets, we trained a classifier to automatically detect the major class of each single cell, as well as classes representing doublets. We first manually annotated clusters to indicate major classes of cells: Neurons, Oligodendrocytes, Astrocytes, Bergman glia, Olfactory ensheathing cells, Satellite glia, Schwann cells, Ependymal, Choroid, Immune, and Vascular. For some of these classes, we distinguished proliferating cells (e.g. Cycling oligodendrocytes, i.e. OPCs). We also manually identified clusters that were clearly doublets between these major classes (e.g. Vascular-Neurons) as well as clusters that were of poor quality.

We then trained a support vector classifier to discriminate all of these labels, using the training set of preliminary clusters manually annotated with class labels. We sampled 100 cells per cluster and used 80% of this dataset to optimize the classifier, and the remaining 20% to assess performance. On average, the classification accuracy was 93% for non-cycling cells. The precision and recall for neurons was 93% and 99%, respectively. That is, 99% of all neurons were classified correctly, and 93% of all cells classified as neurons were actually neurons. The classifier struggled to distinguish cycling cells, presumably because they shared most gene expression with their non-cycling counterparts. For this reason, we always pooled cycling and non-cycling cells after classification. The table below shows the accuracy for all major classes of interest:

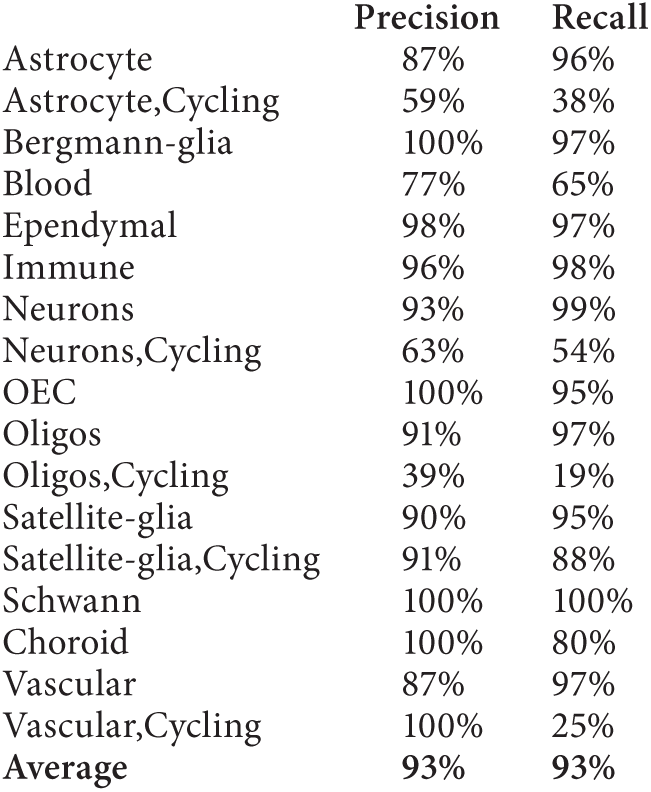

|  | Precision | Recall |
| --- | --- | --- |
| Astrocyte | 87% | 96% |
| Astrocyte,Cycling | 59% | 38% |
| Bergmann-glia | 100% | 97% |
| Blood | 77% | 65% |
| Ependymal | 98% | 97% |
| Immune | 96% | 98% |
| Neurons | 93% | 99% |
| Neurons,Cycling | 63% | 54% |
| OEC | 100% | 95% |
| Oligos | 91% | 97% |
| Oligos,Cycling | 39% | 19% |
| Satellite-glia | 90% | 95% |
| Satellite-glia,Cycling | 91% | 88% |
| Schwann | 100% | 100% |
| Choroid | 100% | 80% |
| Vascular | 87% | 97% |
| Vascular,Cycling | 100% | 25% |
| Average | 93% | 93% |

We used this classifier to individually assess the class identity of each cell in each dataset, and to pool cells by major class into new files (with neurons further separated by tissue).

### Removing doublets

We expected about 2% of all cells to be doublets. Preliminary exploratory analysis (including by generating simulated doublets) showed that most doublets would either form separate clusters, or would tend to end up at the fringes of other clusters (in graph embeddings, and in t-SNE). To eliminate many doublets, we (1) removed clusters classified with ambiguous labels; (2) removed cells classified with a different label from the majority of cells in its clusters; (3) removed outliers when clustering, typically on the fringes of clusters in t-SNE space.

#### Level 1 analysis

We pooled samples by tissue and performed manifold learning, clustering, classification, gene enrichment, and marker gene detection (see below for details on these procedures).

#### Level 2 analysis

We split cells by major class according to the class assignment probability. For each cluster at level 1, we removed cells with conflicting classification (i.e. cells classified as Neuron in a cluster where the majority of cells are classified as Vascular). We performed the same analysis steps as for Level 1.

#### Level 3 analysis (neurons)

Because of the way we had dissected the brain, we would expect some clusters to appear in multiple tissues. For example, our olfactory sample included the anterior-most part of the cortex and underlying tissue (intended to cover the anterior olfactory nucleus), and could overlap with the cortex samples. In order to allow clusters to merge across such boundaries, and in order to improve resolution in clustering, we pooled cells in broader categories, and split them by (mostly) neurotransmitter, as follows:

| Region | Class |
| --- | --- |
| Spinal cord | GABAergic, glycinergic |
| Spinal cord | Glutamatergic |
| PNS | All |
| Hypothalamus | Peptidergic |
| Hindbrain | GABAergic, glycinergic |
| Hindbrain | Glutamatergic |
| Whole brain | Neuroblasts |
| Forebrain | GABAergic |
| Forebrain | Glutamatergic |
| Di- and mesencephalon | GABAergic |
| Di- and mesencephalon | Glutamatergic |
| Whole brain | Granule cells |
| Whole brain | Cholinergic and monoaminergic |
| Striatum | Medium spiny neurons |

Note that we pooled granule cells of the dentate gyrus and the cerebellum not because we think they are related (they are not), but because they are both extremely abundant and tended to skew manifold learning when included with other cells.

#### Level 4 analysis

Despite our efforts, at level 3 there remained still some clusters that were suspected doublets, as well as over-split clusters that lacked clearly defining gene expression differences. We therefore manually curated all clusters, merging some and eliminating others. We then recomputed the manifolds, but did not recluster.

#### Level 5 analysis

To create the final consolidated dataset, we extensively annotated and named each cluster (Table S3). We pooled all cells into a single file along with all metadata and annotations, and performed gene enrichment analysis and marker gene set discovery on this dataset. The level 5 analysis was the basis for all downstream analysis.

#### Level 6 analysis

Finally, level 6 is identical to level 5, but organized into subsets accroding to the taxonomy (Fig. S3). This provides gene enrichment analysis and marker gene set discovery, individually for each taxon.

### Manifold learning

Each individual cell can be viewed as a point in a high-dimensional space, with coordinates given by the expression of every gene. This space would have about 27,000 dimensions, one per gene. In principle, cell types can be viewed as high-density regions in this space, and clustering methods can be used to find them.

In some sense, cells reside on a low-dimensional manifold in the high-dimensional gene expression space. However, the high dimensionality and sparseness of this space creates the “curse of dimensionality”, where distance measures essentially stop making sense. A second issue concerns measurement noise, with generally low counts and large numbers of dropouts (false negatives). Both of these issues can be mitigated by (1) selecting a reduced set of informative genes and (2) linearly projecting the data to a transformed space where each coordinate corresponds to many co-regulated genes. The most effective way of selecting informative genes, would be to select them relative to known classes. We therefore developed a staged procedure to learn the manifold.

We first selected 1000 informative genes by fitting a support-vector regression to the coefficient of variation (CV) as a function of the mean, and selecting genes having the greatest offset from the fitted curve; this would correspond to genes with higher-than-expected variance. We normalized each cell to a sum of 5,000 molecules (UMIs), then log-transformed and subtracted the mean (per gene).

We then used principal component analysis (PCA) to both reduce noise and to reduce the gene expression space further. Dropping non-significant principal components (Kolmogorov-Smirnov test, *P* < 0.05) reduced the space to a few tens of dimensions (typically about forty).

Given a reduced and denoised dataset, we next sought to learn the shape of the manifold of cells (that is, the underlying lower-dimensional gene expression space on which cells are preferentially located). Examining the PCA revealed that the manifold consisted of feather-like, elongated structures, extending variously into the different principal components. We found that the manifold was structured at many levels, ranging from broadly different classes of cells, individual cell types, to more subtle sub-types or states.

We constructed a balanced mutual k nearest-neighbor (KNN) graph with k = 100 using Euclidean distance in the space of significant components. We allowed a maximum of 200 incoming edges to each cell and then dropped all non-mutual edges. We performed Jaccard multilevel community clustering on this graph to define a preliminary set of cell types/states.

Given this preliminary clustering, we were able to select an even more informative set of 500 genes, by calculating an enrichment score (see below) for each cluster, and selecting the *500/n_clusters* most highly enriched genes for each cluster.

Next, we repeated the procedure (PCA, mutual KNN, clustering) with modifications as follows. First, for computing the PCA transform, we limited the number of cells from the largest clusters to contribute max 20% of the total cells (to avoid skewing the PCA towards dominant cell types; note that we still kept all cells in the dataset, only masking those cells when computing the PCA transformation matrix). Second, we computed a balanced KNN as before but we assigned weights *w*(*i,j*)=1/*k*^*α*, where *k* is the rank of *j* among the neighbors of *i* and *a* is a power that sets the scale of the weights. Large values of a will emphasize local neighborhoods, whereas smaller values will emphasize global structure, but in both cases, both local and global structures are accounted for. For practical purposes, we calculated the multiscale graph only up to *k* = 100 (beyond which the edge weights are vanishing), and we used *a* = 1. Using a fixed maximal *k* also ensured that the algorithm remained linear in the number of cells. We call this a multiscale KNN, and stored both the KNN and the mutual KNN for use in further clustering and visualization (available as column graphs named KNN and MKNN in the Loom files).

We projected the multiscale KNN graph to two dimensions using a modified t-SNE algorithm we call graph t-SNE (gt-SNE). In contrast to standard t-SNE, which is based on distance measures, we directly projected the multiscale KNN, which is based on multiscale weighted ranks. We achieved this by replacing the distance matrix P in regular t-SNE with the distance matrix of the weighted multiscale KNN graph. The result was a more accurate projection of the graph itself, with more compact and well-defined neighborhoods. We stored the gt-SNE embedding as column attributes `_X` and `_Y` in the Loom files.

### Clustering

We performed clustering on the multiscale KNN graph. We used Louvain multilevel community clustering (Blondel et al., 2008). However, modularity-based graph clustering suffers a well-known resolution limit (Fortunato and Barthélemy, 2007), failing to find small clusters even when they are perfectly unambiguously defined. Some variants (so called resolution limit-free algorithms) can be tuned to detect smaller clusters, but at the expense of breaking up large clusters. To circumvent this issue, we exploited the fact that we had both a graph, and an embedding of the graph in two dimensions. We first used Louvain clustering on the graph to find most clusters, and then isolated and re-clustered each cluster using DBSCAN in the low-dimensional space. We call this approach “Polished Louvain”.

In more detail, we first performed Louvain community detection on the MKNN graph, with resolution set to 1.0 (except for level 3 where we used 0.6 for astrocytes, 0.35 for sensory neurons and 0.6 for granule cells).

We marked cells as outliers if they (1) belonged to clusters with less than ten cells; or (2) were marked as outliers by DBSCAN (on the 2D embedding) with ε set to the 80th percentile of the distance to the kth nearest neighbor and min_samples = 10; or (3) if more than 80% of the cell’s nearest neighbors belonged to a different cluster.

Next, we isolated each cluster and considered it for further splitting, in the 2D space of the gt-SNE embedding. We centered it using PCA and standardized it by subtracting the mean and dividing by the standard deviation. We marked the cluster for splitting if it now showed three or more outliers based on the median absolute deviation (MAD) with threshold 3.5. We also marked the cluster for splitting if more than 5% of the cells (or 25 cells, whichever is larger) were located at a distance greater than the 70th percentile of the distance to the kth nearest neighbor.

If a cluster was marked for splitting, we performed DBSCAN on that cluster with ε set to the 70th percentile of the distance to the kth nearest neighbor and min_samples = 5% of the cells (or 25 cells, whichever is larger).

Finally, we set the cluster label of each cell to the majority label of its ten nearest neighbors. We stored cluster labels as column attribute Clusters in the Loom files (integer ranging from 0 to *n*). At level 5 and 6, `cluster` names are given by the column attribute `ClusterName`.

### Gene enrichment

To aid interpretation of the data (and for gene selection, as noted above), we computed a set of genes enriched in each cluster. We computed an enrichment statistic *E*_*i,j*_ for gene *i* and cluster *j*, as follows:

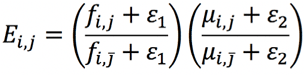

where *f* is the fraction of non-zero expression values, and *μ* is the mean expression, and *j* with overdash denotes cells not in the cluster. Small constants *ε*_1_ = 0.1 and *ε*_2_ = 0.01 are added to prevent the enrichment score from going to infinity as the mean or non-zero fractions go to zero. Enrichment scores are available as matrix layer `enrichment` in the aggregated Loom files (named “…agg.loom”). We also computed an enrichment *q* value by shuffling the expression matrix, available as layer `enrichment_q`. To find genes enriched at a 10% false discovery rate, for example, simply select genes with *q* scores below 0.1.

### Trinarization

It is often useful to estimate (for each cluster) if a gene is likely expressed, not expressed, or we are not sure. That is, we want to trinarize the raw expression data into calls of expressed, not expressed, and indeterminate. Here we used a Bayesian beta-binomial model to trinarize the raw data.

The model applies to a cluster of cells representing a putatively homogeneous population. In this cluster, we have measured gene expression in n cells, and for each cell we have either detected the gene, or not. Given detection in *k* out of *n* cells, we want to know the underlying population frequency of expression, *Θ*. The observed fraction of expressing cells can be expressed conditional on the number of cells and the population expression frequency. By providing a prior on *Θ*, we can derive the posterior distribution of *Θ* given the observed number of detections:

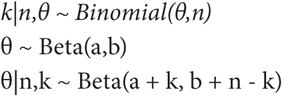

The Beta distribution is the conjugate prior to the Binomial, and as a consequence the posterior distribution is also Beta, and can be calculated simply by updating the parameters. Setting *a* = *b* = 1 results in a non-informative uniform prior. Here, we used instead a weakly informative prior with *a* = 1.5, *b* = 2, which slightly favours the “not expressed” and “indeterminate” calls.

Using this model to trinarize gene expression, we call a gene expressed when *P(Θ > f) > (1 - PEP)*, where f is the population fraction of cells expressing the gene, and *PEP* is the desired posterior error probability (also called local false discovery rate). For example, with *PEP = 0.05*, there is less than 5% risk, given the observations, that the expressed call is wrong. Similarly, we call a gene not expressed when *P(Θ > f) < PEP*. For values between *1-PEP* and *PEP*, we call the gene indeterminate. Note that *PEP* is applied individually to each gene (hence, “local FDR”) and the actual genome-wide FDR will be strictly equal to or lower than *PEP*.

The probability *P(Θ > f)* can be calculated as:

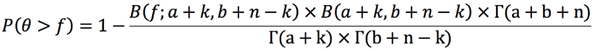

The formula was derived by evaluating the expression Probability[x>f, {x~BetaDistribution[a+k, b+n-k]}] in Mathematica (version 10, Wolfram Research Inc.). Here, *B(z; a, b)* is the regularized incomplete beta function, *B(a, b)* is the beta function, and *Γ* is the gamma function.

Evaluating this function, for a given *k* and *n* (and hyperparameters *f*, *a* and *b*) yields a probability *P*, which we compare to the thresholds *1-PEP* and *PEP* to give the gene an expression call. We used *a=1.5*, *b=2*, *f=0.2* and *PEP=0.05* to make the calls in this paper, unless otherwise indicated. Thus a gene was considered expressed if it was estimated to be present in at least 20% of the cells with no more than 5% posterior error probability.

Note that the formula as written suffers from numerical instability when evaluated at finite precision. This problem can be avoided by using logarithms of the beta and gamma functions, and then exponentiating. See the source code of function `p_half` in the file `diff_exp.py` for a complete, numerically stable implementation.

Trinarization scores are available in layer `trinaries` in the aggregated loom files.

### Marker gene set discovery

Many, even most, of the cell types described in this paper were not previously associated with known makers. We therefore designed an algorithm to automatically propose marker sets for all clusters. Here, we define a marker gene set as a set of genes that are all expressed in a given cluster, but not all expressed in any other cluster. We used trinarization to judge if a gene is expressed or not in each cluster.

Given a cluster, we first selected the most highly enriched gene, which would often not be unique to that cluster, but highly selective for a small number of closely related clusters. Next, we added the most specific gene, based on trinarization with a *PEP* of 0.05. This gene was very often specific to a very small number of clusters, and using the first two genes together would often lead to fully specific marker combinations. However, sometimes adding more genes would be necessary.

We added genes one at a time by picking the most selective gene, in combination with the previously selected genes. When more than one gene was equally selective, we picked the one that was most highly enriched. We defined selectivity as the reciprocal of the number of clusters that would be selected given the current gene set and the trinarization scores. That is, gene set that would be all-positive in *k* clusters would have selectivity *1/k*. Adding more genes rapidly drove selectivity towards 1.

We generated gene sets in this manner for all clusters, with up to six genes per cluster. We also calculated the cumulative selectivity, specificity (difference between the posterior probability for the best cluster and that of the second-best cluster), and robustness (the posterior probability that all genes would be detected in the cluster, based on trinarization scores). We reported these statistics cumulatively for *n* = 1, 2, 3, 4, 5 and 6 genes. Generally, robustness drops as more genes are added, while selectivity increases. Specificity tends to increase as the gene set becomes more selective, but then decrease as it becomes less robust.

We note that marker gene sets are excellent candidates to use for experimentally identifying cell types, e.g. based on genetic or antibody labelling. Marker gene sets and associated statistics for all clusters are provided in the wiki, and in the Loom files under column attributes `MarkerGenes, MarkerSelectivity, MarkerSpecificity and MarkerRobustness`.

### Dendrogram construction

All linkage and distance calculations were performed after *log*_2_(*x*+1) transformation. The starting point of the dendrogram construction was the 265 clusters. For each gene, we computed average expression, trinarization with *f* = 0.2, trinarization with *f* = 0.05 and enrichment score. For each cluster we also know the number of cells, annotations, tissue distribution and samples of origin.

We defined major classes of cell types based on prior knowledge: neurons, astroependymal, oligodendrocytes, vascular (without VLMC), immune cells and neural crest-like. For each class, we defined pan-enriched genes based on the trinarization 5% score. Each class (except neurons) was tested against neurons, to find all the genes where the fraction of clusters with trinarization score = 1 in the class was greater than the fraction of clusters with trinarization score > 0.9 among neurons.

In order to suppress batch effects (mainly due to ambient oligodenderocyte RNA in hindbrain and spinal cord samples), we collected the unique set of genes pan-enriched in the non-neuronal clusters, as well as a set of non-neuronal genes that we believe to have tendency to appear in floating RNA (*Trf*, *Plp1*, *Mog*, *Mobp*, *Mfge8*, *Mbp*, *Hbb-bs*, *H2-DMb2*) and a set of immediate early genes (*Fos*, *Jun*, *Junb*, *Egr1*). These genes were set to zero within the neuronal clusters to avoid any batch effect when clustering the neuronal clusters. We further removed sex specific genes (*Xist*, *Tsix*, *Eif2s3y*, *Ddx3y*, *Uty*, and *Kdm5d*) and immediate early genes *Egr1* and *Jun* from all clusters.

We bounded the number of detected genes in each cluster to the top 5000 genes expressed, followed by scaling the total sum of each cluster profile to 10,000.

Next, we selected genes for linkage analysis: from each cluster select the top *N=28* enriched genes (based on pre-calculated enrichment score), perform initial clustering using linkage (Euclidean distance, Ward in Matlab), and cut the tree based on distance criterion 50. This clustering aimed to capture the coarse structure of the hierarchy. For each of the resulting clusters, we calculated the enrichment score as the mean over the cluster divided by the total sum and selected the *1.5N* top genes. These were added to the previously selected genes.

Finally, we built the dendrogram using linkage (correlation distance and Ward method).

### Test for dendrogram stability

We tested the stability of the dendrogram structure while changing the number of genes selected for calculating the dendrogram. We selected N in the range 10-44. For each N we repeated the procedure above and stored the selected genes and dendrogram structure. We then examined all branches (junctions) of the reference tree (*N=28*) and compared them to the corresponding branch in the test tree. We derived two stability criteria (1) branches with leafs below having 90% overlap in test compared to reference, (2) branches with exactly the same set of clusters and the same order of the leafs. For each branch we calculated the fraction of cases that either criteria (1) or (2) occurred. More than 65% of the 264 branches had probability of 1 and 94% had probability greater than 0.5 based on criterion (1). Based on the more stringent criterion (2) more than 50% had probability of 1 and about 85% greater than 0.5.

### Testing for dendrogram without any gene exclusion

In the dendrogram construction described above we used several steps of exclusion genes either from all clusters or from the neuronal clusters in particular. This was done due to our observation of background levels of gene detection which seemed to be depend on very abundant cell types the dissected region (e.g. oligodendrocytes in hindbrain or enteric glia in the enteric nervous system). This is likely because of floating RNA coming from dead cells or doublets either with abundant cells or parts of broken cells. Still, due to the risk of misinterpreting the data we also constructed the dendrogram based on similar procedure but without any gene exclusion. The resulting dendrogram was not fundamentally changed from Fig. 1C, but included a few key differences which we believe are technical artefacts. First, enteric neurons clustered together with the enteric glia probably due to fact that enteric glia were extremely abundant in the tissue. This created a big enough change that the other PNS neurons created a separate branch disconnected from the other neurons. Second, the olfactory bulb inhibitory neurons were placed next to the MSNs. This branch in turn was connected to a branch mainly containing neuroblasts. Finally, the OPC cluster was placed next to the SZNBL cluster probably because of strong cell-cycle signal.

### Spatial correlation analysis

Our aim here is to try and map the gene expression profile at the cluster level to the in-situ hybridization atlas of the Allen Institute for Brain Research. The Allen Mouse Brain Atlas was summarized into a 200 µm voxel dataset, providing the gene expression profile (all genes) for each voxel. In this analysis we used simple correlation between the voxel gene expression (from in situ hybridization) and the cluster gene expression profile (from scRNAseq).

For each gene, the voxel data is a 67×41×58 (rows × columns × depth) array, giving an “energy” value representing the expression. In addition, for each voxel we know the anatomical annotation. The Allen Brain reference atlas is given at a finer resolution with voxels of 25µm (528×320×456). In order to achieve finer resolution and smoother images we used linear interpolation of the coarse (200 µm) in-situ data into the finer grid (25 µm). For annotation we used the color code of the Allen reference atlas.

Since many genes have information only from sagittal sections of one hemisphere, we can neglect one hemisphere also from the genes that have coronal data. Coronal data is preferred since it has better sampling.

### Procedure

First, we define the energy of any voxel outside the valid domain to −1. We define genes as high-quality

